# The functional type VI secretion system of *Sinorhizobium fredii* USDA257 is required for a successful nodulation with *Glycine max* cv Pekin

**DOI:** 10.1101/2023.12.14.571490

**Authors:** Pedro José Reyes-Pérez, Ana Sánchez-Reina, Cristina Civantos, Natalia Moreno-de Castro, Francisco Javier Ollero, Jacinto Gandullo, Irene Jiménez-Guerrero, Patricia Bernal, Francisco Pérez-Montaño

**Affiliations:** 1. Departamento de Microbiología, Facultad de Biología, Universidad de Sevilla, Avda. Reina Mercedes 6, Sevilla, Spain; 2. Departamento de Biología Vegetal y Ecología, Facultad de Biología, Universidad de Sevilla, Avda. Reina Mercedes 6, Sevilla, Spain

**Author notes:** Address correspondence to Irene Jiménez-Guerrero, Patricia Bernal and Francisco Pérez- Montaño, and.

**Keywords:** rhizobium, legume, symbiosis, nodulation, competition, type VI secretion system, effector

## Abstract

For agriculture, the symbiosis carried out by rhizobia with legumes stands out as crucial for both economic and environmental reasons. In this process, the bacteria colonize the roots of the plants, inducing the formation of plant organs called nodules. Within these structures, rhizobia fix the environmental nitrogen into ammonia reducing the demand for this essential element required for plant growth. Various bacterial secretion systems (TXSS, Type X Secretion System) are involved in the establishment of this symbiosis, with the T3SS being the most extensively studied. The T6SS is a nanoweapon present in 25% of gram-negative bacteria, commonly used against other gram-negative bacteria, though some of them use it to manipulate eukaryotic cells.

Interestingly, although T6SS is widely distributed among rhizobia, whether it has a specific role in symbiosis with legumes remains elusive. *Sinorhizobium fredii* USDA257 is a fast-growing rhizobium with the capacity to nodulate a great variety of legume plants. This strain harbors a single T6SS cluster, containing the genes encoding all the structural components of the system and two genes encoding potential effectors that could target the cell wall of the plants and/or be acting as a toxin/antitoxin system. We have demonstrated that this system is active and can be induced in poor culture media. In addition, we have seen by fluorescence microscopy that the T6SS is active in nodules. Competition assays between USDA257 and different preys have shown that USDA257 cannot kill any of them using its T6SS under tested conditions. By constract, nodulation assays demonstrated that USDA257 utilizes this protein secretion system to enhance nodulation and competitiveness with its host *Glycine max* cv Pekin.

## 1. Introduction

Rhizobia are α- and ß-Proteobacteria soil-borne microorganisms frequently found on leguminous plant roots and rhizosphere. This environment is highly appropriate for rhizobia development and, eventually, they form a symbiotic relationship with the leguminous plant. The rhizosphere provides rhizobia with protection from desiccation, extreme temperatures and light stress. At the same time, the legume supplies the bacteria with nutrients exuded from the roots, including amino acids, organic acids, sugars, aromatic compounds and secondary metabolites (Walker *et al*., 2003). The flavonoids are secondary metabolites exudated by the plants, that initiate the molecular dialogue between rhizobia and legume. This dialogue culminates with the formation of new organs in roots called nodules, where the bacterial reduction of atmospheric nitrogen to the most-needed form -ammonia- takes place (Oldroyd & Downie, 2008). Thus, when recognized by the appropriate rhizobia, the flavonoids induce the production of rhizobial signal molecules, called nodulation factors (Nod factors). These signal molecules can be specifically recognized by legume plants, initiating the nodulation process that will culminate with nodule development, its occupation and the transformation of the rhizobium into a nitrogen-fixing cell, the bacteroid (Oldroyd *et al*., 2011). Besides the molecular recognition mediated by flavonoids and Nod factors, the success of the nodulation process also depends on secretion systems effectors. Thus, in the last fifteen years, the effector proteins secreted through the rhizobial type III secretion system (T3SS) have been proven important, and in some cases essential, for the symbiotic performance of several rhizobial genera, such as *Sinorhizhobium*, *Rhizobium*, *Bradyrhizobium* and *Mesorhizobium* (López-Baena *et al*., 2016; Jiménez- Guerrero *et al*., 2021, 2022). A novel secretion system has been described in the last two decades, the type VI secretion system (T6SS). In fact, this machinery was first identified in *Rhizobium leguminosarum* (Bladergroen *et al*., 2003), although the term was not stablished until 2006, when the T6SS of *Vibrio cholerae* and *Pseudomonas aeruginosa* were simultaneously characterized (Mougous, 2006; Pukatzki *et al*., 2009). The T6SS is present in 25% of gram-negative bacteria, mainly in the Proteobacteria phylum, where the α-, β- and γ-proteobacteria classes are included (Boyer *et al*., 2009). Generally, this system secretes effectors/toxins into prokaryotic cells, playing a critical role in inter-bacterial competition (Ho, *et al*., 2014). However, some type VI-secreted effectors (T6Es) can target eukaryotic organism cells, having the ability to manipulate the host during an infective process (Hachani *et al*., 2016).

The specific role of the T6Es in rhizobia symbiosis is mostly unknown, although recent works suggest that these proteins could exert either neutral, positive, or negative effects, depending on the symbiotic pair. Thus, it has been reported that the T6SS of *Paraburkholderia phymatum* STM 815^T^ and *Azorhizobium caulinodans* do not appear to be directly implicated in the symbiotic effectiveness with *Vigna unguiculata* and *Sesbania rostrata*, respectively. However, it is described that both systems are involved in symbiotic competitiveness against rhizobial competitors for nodulation (de Campos *et al*., 2017; Lin *et al*., 2018). Conversely, in both *Rhizobium etli* Mim1 and *Bradyrhizobium* sp. LmicA16, the T6SS is required for efficient nodulation with *Phaseolus vulgaris* and *Lupinus* spp, respectively (Salinero-Lanzarote *et al*., 2019; Tighilt *et al*., 2022). In case of *R. etli* Mim1, this system is synthesized at high cell densities, in the presence of root exudates and within host-plant nodules (Salinero- Lanzarote *et al*., 2019), being the T6E Re78 beneficial for competitiveness in bean nodule occupancy (de Souza *et al*., 2023). Finally, *Rhizobium leguminosarum* RBL5787 is unable to form nitrogen-fixing nodules on pea (*Pisum sativum*) due to the presence of a functional T6SS (Bladergroen *et al*., 2003).

Structurally, the T6SS is a multiprotein complex composed of thirteen main constituents. The genes encoding these proteins are grouped in genetic clusters and named *tss* (type <u>s</u>ix <u>s</u>ecretion) (Ho *et al*., 2014). In some cases, there is an additional set of genes, named *tag* genes (type <u>s</u>ix <u>a</u>ccessory genes), that encode for accessory proteins with complementary functions . Those proteins have been involved in regulation (TagF, TagQ, TagR, TagS, TagT) (Lin *et al*., 2018; Hsu *et al*., 2009) and fine-tuning the system assembly (TagP, TagL, TagN, TagW, TagA, TagB, TagJ) (Aschtgen *et al*., 2010; Santin *et al*., 2018; Bernal *et al*., 2021). The T6SS is structured into three main compartments, and the TssA protein orchestrates the assembly among the different parts. The membrane complex, formed by TssJ, TssL and TssM, anchors the system to the cell envelope. TssA interacts with the membrane complex and recruits the baseplate, composed by TssK, TssE, TssF and TssG. In this new position, TssA primes polymerization of the tail, formed by an inner tube (Hcp), surrounded by a contractile sheath (TssB and TssC) and ended in a needle-shaped tip (VrgG and PAAR). The T6Es can be transported inside the tube, or connected to the tip, being released into the intracellular environment of the target cell upon sheath contraction. The sheath, which remains in the cytosol, is then disassembled by an AAA^+^ ATPase, named ClpV (or TssH), which plays a crucial role in the recycling of the system (Cherrak *et al*., 2018).

Genes encoding effector/toxin proteins can be found within the T6SS cluster or scattered around the genome. These genes are usually located downstream genes encoding VgrG, Hcp or/and PAAR proteins. In some cases, T6Es are encoded by the same gene that encodes for Hcp, VrgG or PAAR proteins and, therefore, they are fused to them at the C-terminal end. Consequently, structural proteins with a C-terminal cytotoxic domain are termed evolved, named as Hcp_e_, VrgG_e_ or PAAR_e_ (Durand *et al*., 2014). The antibacterial T6Es can target the periplasm; these include peptidoglycan hydrolases (Tae1-4 and Tge1-3), phospholipases (Tle1-4) and pore-forming effectors (VasX). Other T6Es act in the bacterial cytoplasm, such as the nucleases effectors (Tde1 and Tke2), or those that degrade essential cofactors - NAD(P)^+^, like the hydrolase effectors (Tse6/Tne1 and Tne2) (Coulthurst, 2019). Bacteria with a functional T6SS produce immunity proteins to protect from sister cells attack and self-intoxication. Contrary, T6Es delivered into eukaryotic host cells are less widespread than antimicrobial effectors, but the few identified so far are involved in different steps of host manipulation to promote bacterial infection (Hachani *et al*., 2016).

*Sinorhizobium fredii* USDA257, hereafter USDA257, is a fast-growing rhizobium that was isolated from wild soybean (*Glycine soja*), but is also able to form nitrogen-fixing nodules in a wide variety of legume species, such as *G. max*, *S. rostrata, V. radiata, P. vulgaris*, *Cajanus cajan*, *Lotus japonicus*, and *L. burtii,* among others (Pueppke & Broughton, 1999). USDA257 is one of the most versatile rhizobia, along with other bacteria of the same species, *S. fredii* NGR234 and *S. fredii* HH103. Interestingly, only USDA257 presents both T3SS and T6SS machinery. The T3SS of USDA257 has been extensively studied,playing a prominent role in symbiosis, as well asin tdetermining of its host-range of nodulation (Staehelin & Krishnan, 2015). Legume recognition of type III-secreted effectors(T3Es) can exert both positive (induction of nodulation) or negative (inhibition of nodulation) extreme effects, depending on the plant cultivar (Jiménez-Guerrero *et al*., 2022). However, ecological and physiological function of the USDA257 T6SS is completely unknown in the *Sinorhizobium* genera.

In this work, we identified and characterized the USDA257 T6SS, which exhibits novel characteristics of the structural proteins, implying an apparatus with a distinctive assembly and a definite set of T6Es. We showed that this USDA257 secretion system is especially active in non-rich media during stationary phase of growth and when colonizes legume root nodules. In fact, nodulation assays demonstrated that USDA257 utilizes this protein secretion system to enhance nodulation and competitiveness with *G. max* cv Pekin. According to *in silico* predictions, improvement in the symbiotic performance could be directly mediated by essential T6Es and/or indirectly by secreting antibacterial toxins during the nodulation process.

## 2. Materials and Methods

### 2.1. Bioinformatical analysis

Sequences of 161 TssB proteins from 153 strains, belonging to 12 genera were obtained and compiled by using the BLASTp tool from NCBI website (Boratyn *et al*., 2013). Sequences were aligned with the ClustalW software (Sievers *et al*., 2011) and the construction of the phylogenetic tree was carried out using the MEGA7 software (Kumar *et al*., 2016), applying the Maximum Likehood algorithm (Le *et al*., 2008) and a JTT matrix model (bootstrap value=500). The phylogenetic tree was customized with the iTOL tool (Letunic & Bork, 2016). Amino acid sequence searches were carried out using SMART (Letunic *et al*., 2015) and Pfam (Finn *et al*., 2016). The Protein Homology/analogy Recognition Engine (Phyre^2^) server was used to perform a template-based approach to predict protein structural homology (Kelley *et al*., 2015). An additional analysis to identify structural homologs was performed using the recent algorithm Foldseek (Van Kempen *et al*., 2024), that is a structural alignment tool based on a structural protein alphabet of tertiary interactions that operates using predicted models from the Alphafold database (Abramson *et al*., 2024). Molecular weight and isoelectric point for each protein were calculated with the ExPaSy software (Artimo *et al*., 2012). PSORTb server was used to predict sub-cellular location of proteins (Yu *et al*., 2010), TMHMM software to predict transmembrane domains (Krogh *et al*., 2001), and SignalP and PSORTb to predict signal peptides (Almagro Armenteros *et al*., 2019). The figures for the cluster alignments were created using Easyfig (Sullivan *et al*., 2011).

### 2.2. Bacterial strains and plasmids

Bacterial strains and plasmids used in this work are listed in **Table S1**. Rhizobial strains used in this study were grown at 28 °C on tryptone yeast (TY) medium (Beringer, 1974), yeast extract mannitol (YM) medium (Vincent, 1970) or minimal (MM) medium (Robertsen *et al*., 1981) with two different mannitol concentrations (3 or 10 g mL^-1^, YM3/MM3 or YM10/MM10, respectively). *Agrobacterium tumefaciens* and *Pectobacterium carotovorum* strains were cultured on LB medium (Bertani, 1951) at 37°C in the first case, and 28 °C in the last two ones. When required, the media were supplemented with the appropriate antibiotics, as previously described (Lamrabet *et al*., 1999). Genistein was dissolved in ethanol and used at 1 µg ml^-1^ to give a final concentration of 3.7 µM.

Plasmids were transferred from *E. coli* to *Sinorhizobium* strains by triparental conjugation, as described by Simon (1984), using plasmid pRK2013 as helper (Figurski & Helinski, 1979). Recombinant DNA techniques were performed according to the general protocols of Sambrook *et al*. (1989). PCR amplifications were performed as previously described López-Baena *et al*., (2009). Primer pairs used for the amplification of the USDA257 genes are summarized in **Table S2**.

For the obtention of a USDA257 *tssA* mutant, an internal fragment of this gene was amplified using specific primers, digested with *Eco*RI and *Bam*HI restriction enzymes (both restriction sites were added in forward and reverse 5′ primer ends, respectively) and cloned into pK18*mob* vector, which was previously digested with the same enzymes, obtaining plasmid pMUS1480. This construction was employed for the homogenotization of the mutated version of *tssA* in USDA257, generating the mutant strain. Double recombination events was confirmed by Southern blot, PCR and sequencing (data not shown). For hybridization, DNA was blotted to Hybond-N nylon membranes (Amersham, UK), and the DigDNA method of Roche (Switzerland) was employed according to the manufacturer’s instructions (**Fig. S1**).

Promotor regions of the *ppkA* USDA257 T6SS gene was amplified with specific primers (575 bp) and the resulting DNA fragment was digested with *Eco*RI and *Xba*I (both restriction sites were added in forward and reverse 5′ primer ends, respectively) and cloned into plasmid pMP220, upstream of the *lacZ* gene, being previously digested with same enzymes, obtaining plasmid pMP220::P_ppkA._

Construction of the dual reporter strain of USDA257 was performed using the plasmid developed by Shamal and Chatterjee (2021) and inserting the promoter region of the *ppkA* gene (530 pb), that was previously amplified and digested with *Eco*RI (restriction sites were added in forward and reverse 5′ primer ends). Briefly, transcriptional fusion was constructed by fusing the promoter region upstream of the *ppkA* gene (P_ppkA_) to the red fluorescent protein gene (*rfp*) and cloned into a pBBR-MCS-4 plasmid, that is constitutively expressing the *gfp* gene (P_kan_::*gfp*). This plasmid was conjugated with the USDA257 strain for obtaining the dual-reporter strain of USDA257 (pBBR4::P_kan_::*gfp*- P_ppkA_::*rfp*). As a control, the plasmid pBBR4::P_kan_::*gfp*-P_w/o_::*rfp* (Shamal and Chatterjee, 2021), in which the *rfp* gene lacks the promoter region, was also transferred by conjugation to USDA257. All plasmids were stably maintained during infection.

For USDA257 Hcp antibody production, the *hcp* ORF without the stop codon was amplified by PCR with specific primers and cloned into the entry plasmid pDONR207 and then mobilized into pET42 plasmid following the Gateway cloning system technology (Invitrogen, USA). The generated plasmid was transferred to*E. coli* BL21 (DE3) (Studier and Moffatt, 1986) by the heat shock method (Froger and Hall, 2007).

### 2.3. Determination of β-galactosidase activity

To determinate the β-galactosidase activity, the USDA257 wild-type strain was conjugated with the plasmids pMP220::P_ppkA_ and pMP240, that contains transcriptional fusions between the *ppkA* (USDA257) and *nodA* (*R. leguminosarum*) promoters, respectively, both upstream of the *lacZ* gene. The empty plasmid pMP220 was also transferred to USDA257 and used as negative control. Assays of β-galactosidase activity were carried out as described by Zaat *et al* (Zaat *et al*., 1987). Units of β- galactosidase activity were calculated according to Miller (Miller, 1972). The experiments were repeated four times with three technical replicates each time.

### 2.4. RNA extraction and qRT-PCR experiments

Total RNA was isolated using a High Pure RNA Isolation Kit (Roche, Switzerland), according to the manufacturer’s instructions. Verification of the amount and quality of total RNA samples was carried out using a Nanodrop 1000 spectrophotometer (Thermo Scientific, USA) and a Qubit 2.0 Fluorometer (Invitrogen, USA). Four independent total RNA extractions were obtained for each condition. This (DNA-free) RNA was reverse transcribed into cDNA by using PrimeScript RT reagent Kit with gDNA Eraser (Takara, Japan). Quantitative PCR was performed using a LightCycler 480 (Roche, Switzerland) equipment with the following conditions: 95 °C, 10 min; 95 °C, 30 s; 50 °C, 30 s; 72 °C, 20 s; forty cycles, followed by the melting curve profile from 60 to 95 °C to verify the specificity of the reaction. The USDA257 16S rRNA gene was used as an internal control to normalize gene expression. The fold-changes of four biological samples with three technical replicates of each condition were obtained using the ΔΔCt method (Pfaffl, 2001). Selected genes and primers are listed in **Table S2**.

### 2.5. Expression and purification of recombinant Hcp for specific antibody generation

The *E. coli* BL21(DE3) strain carrying plasmid pET42::*hcp* was inoculated in 5 mL of LB supplemented with carbenicillin and incubated overnight at 37 °C with shaking at 200 rpm. The culture was transferred to 200 mL in the same medium and incubated in the same conditions. When the OD_600_ reached 0.6, protein expression was induced with 1 mM isopropyl β-D-1-thiogalactopyranoside (IPTG). Then, cultures were grown for 4 hours at 37 °C with shaking at 200 rpm. Cells were harvested by centrifugation (5.000 g, 20 min, 4 °C) and the pellet was resuspended in 50 mM Tris-HCl (pH 7.5), 250 mM NaCl (buffer NPI) containing 10 mM imidazole, 1 mg/mL of lysozyme and protease inhibitors (Roche, Switzerland). The suspension was incubated for 30 min and sonicated on ice 5 times for 30 sec, with 30-sec cooling intervals between sonication treatments. Cell debris was eliminated by centrifugation (10.000 g, 30 min, 4 °C). Clarified lysate was filtered with a 0.45 nm filter and incubated in a column containing 2.5 mL of nickel-Sepharose resin (Protino Ni-NTA, Macherey-Nagel, Duren, Germany), previosuly equilibrated with 10 volumes of buffer NPI allowing binding of His-tagged Hcp to Protino Ni-NTA agarose. Columns were washed with 10 volumes of buffer NPI containing 100 mM imidazole and, finally, the His-tagged Hcp protein was eluted off the resin with 1 mL of buffer NPI containing 500 mM imidazole. The His-tagged Hcp protein was subsequently washed with PBS and concentrated using Amicon Ultra Centrifugal Filter Unit (Milipore Sigma, Burlington, MA, USA), following the manufacturer’s instructions. Expression and purification of His-tagged Hcp were verified by SDS-PAGE and confirmed by Western blot using His-tag monoclonal antibody. Polyclonal antibody production was carried out by the “Centro de Experimentación Animal Óscar Pintado” from the University of Sevilla (Spain) following the procedure described by Vidal et al. (1980). through their speedy 28-Day program (1 rabbit; 4 protein injections and 2 bleedings).

### 2.6. Purification and analysis of extracellular proteins

Extracellular and intracellular proteins were recovered following the protocol described by Hatachi et al. (2011) with some modification. Briefly, 20 ml of the different rhizobial cultures grown on an orbital shaker (180 rpm) at 28 °C for 48 h (stationary phase, with an adjusted OD_600_ of 1, which corresponds to approximately 10^9^ bacteria ml^-1^) were centrifuged for 20 min at 10.000 g at 4 °C. Bacterial pellets were normalized and added directly to 200 μl of sample buffer (62.5 mM Tris-HCl [pH 6.8], 2% SDS [m/v], 10% glycerol [v/v], 5% β-mercaptoethanol [m/v], and 0.001% bromophenol blue [m/v]). To eliminate any remaining cell in the supernatant, 3 additional sequential centrifugations (20 min, 10.000 g, 4 °C) were performed. A volumen of 1.8 mL from each culture supernatants were collected and precipitated with trichloroacetic (TCA) acid overnight at -20 °C. The mixtures were centrifuged for 30 min at 16.000 g at 4 °C. Dried pellets, previously washed with 90% acetone, were resuspended in sample buffer. For immunostaining, proteins were separated on SDS 20-4% (m/v) polyacrylamide gels (Bio-Rad, USA) and electroblotted to Immun-Blot polyvinylidene difluoride membranes (Bio-Rad) using a Mini Trans-Blot electrophoretic transfer cell (Bio-Rad). Membranes were blocked with TBS containing 2% (m/v) bovine serum albumin (BSA) and then incubated with antibodies raised against the USDA257 Hcp protein diluted 1:1000 in the same solution. Anti-rabbit HRP-linked antibody was used as secondary antibody and reaction results were visualized using CDP Start Detection Reagent kits (GE Healthcare, USA) according to manufacturer’s instructions. Bands were visualized using an ImageQuant LAS 500 imaging workstation instrument (GE Healthcare, USA).

### 2.7. Interbacterial competition assays

*In vitro* competition assays were performed on the USDA257 T6SS-inducing YM3 and MM3 agar (2% w/v) plates, as previously described (Bernal *et al*., 2021). Briefly, overnight bacterial cultures were washed and adjusted to an OD_600_ of 1.0 in sterile PBS,and mixed at a 1:1 ratio (USDA257:prey). *A. tumefaciens*, *P. carotovorum* and *S. fredii* HH103 were used as preys. 100 µl of mixtures were grown on YM3 (for *A. tumefaciens* and *P. carotovorum*) or MM3 (for *S. fredii* HH103) agar plates at 30 °C for 24 or 48 hours and then collected using an inoculating loop and resuspended in sterile PBS. The outcome of the competition was quantified by counting colony forming units (CFUs) using antibiotic selection of the input (time = 0 hours) and output (time = 24-48 hours). *A. tumefaciens* and *P. carotovorum* prey strains harbored the plasmid pRL662, which confers resistance to gentamicin and was used for antibiotic selection, whereas USDA257 was naturally resistant to streptomycin. Three biologically independent experiments were performed.

### 2.8. Plant tests

For the evaluation of the symbiotic phenotypes in nodulation assays, the wild-type and its derivative mutant strains were grown in YM3 medium. Surface-sterilized seeds of *G. max* cv Pekin were pre-germinated and placed in sterilized Leonard jars, containing Fårhaeus N-free solution (Vincent, 1970). Germinated seeds were then inoculated with 1 mL of bacterial culture with an adjusted OD_600_ of 0.6. Growth conditions were 16 h at 26 °C in the light and 8 h and 18 °C in the dark, with 70% humidity. Nodulation parameters were evaluated after 6 weeks. Shoots were dried at 70 °C for 48 h and weighed. Nodulation experiments were performed three times with five replicates for each treatment.

Competition for nodulation experiments (competitiveness) on *G. max* cv Pekin were performed using USDA257 and the non-T6SS-carrying *S. fredii* HH103 rhizobium . These bacteria were grown to 10^9^ cells ml^-1^, and four to five Leonard jar assemblies containing two plant seedlings each were inoculated with 1 ml of a mixture of bacterial competitors in 1:1, 1:10 and 10:1 ratios. Plants were grown for 6 weeks in a plant growth chamber under the growth conditions described above. To identify bacteria occupying the nodules,100 *G. max* cv Pekin nodules from each treatment were surface sterilized by immersing them in 5% [wt/vol] sodium hypochlorite (for 5 min, followed by five washing steps in sterilized distilled water. The effectiveness of the surface-sterilizing treatment was checked by inoculating TY plates with 20-µl aliquots from the last washing step. Individual surface sterilized nodules were crushed in 30 µl of sterilized distilled water, and 20-µl aliquots were used to inoculate TY plates. Nodule occupancy was determined by assessing the ratio of differential natural antibiotic resistant of isolates (Rifampicin for USDA257 and Streptomycin for HH103). At least 10 colonies from each isolate were analyzed in order to check the possibility of nodules containing both inoculants.

For nodule occupancy visualization, 30-day-old *L. burttii* nodules formed in plants inoculated with the dual fluorescent reporters of USDA257 were embedded in 6% agarose in water and sliced in thick layer sections (50 µm) using a Leica VT 1000S vibratome (Wetzlar, Germany). Functional fluorophores of sections of nodules were stained with 0.04% calcofluor and observed by using a Leica Stellaris 8 SPE Confocal Microscope (Leica Microsystems) (Jena, Germany) fluorescence microscope as previously described (Kawarada *et al*., 2017). The imagés contrast and intensity were adjusted using ImageJ software (Schneider *et al*., 2012).

### 2.9. Statistical analysis

The statistical tests performed in this work and indicated in the Figure legends were done with Prism 8 (GraphPad, La Jolla, CA, USA).

## 3. Results

### 3.1. Genome-wide screening for T6SSs in N_2_-fixing bacteria

We have performed a T6SS phylogenetic analyses of 160 N_2_-fixing bacteria from 13 different genera all belonging to the Phylum Proteobacteria and including the main genera containing root-nodule forming bacteria (**Table S3**). The selected species belong to the order *Hyphomicrobiales* (better known as *Rhizobiales*) from the α-Proteobacteria class and the order *Bulkholderiales* from the β-Proteobacteria class. We have included well-described *Agrobacterium* and *Pseudomonas* T6SSs to identify and locate the previously described T6SS phylogenetic groups (Wettstadt *et al*., 2020). The tree displays in **Fig. 1** contains 160 TssB proteins, showing the phylogenetic distribution of selected rhizobia T6SSs. Our analysis shows that rhizobial T6SSs are distributed among the five main clades previously described (Boyer *et al*., 2009). Most of them, 140 (87.5%) belong to groups 3 (54, 33.75%) and 5 (86, 53.75%) (**Fig. 1**). Group 3 contains a great variety of species from the *Ensinfer*, *Rhizobium*, *Bradyrhizobium* and *Mesorhizobium* genera among others, as well as the first-discovered *P. aeruginosa* H1- T6SS. Group 5 contains mostly *Rhizobium* and *Azorhizbium* species and includes the well-studied *A. tumefaciens* T6SS. Although phylogenetically-distance *Cupriavidus* and *Parabulkholderia* species can be found in both groups. Minority groups 2 and 4 principally contain *Paraburkholderia* species (2, 1.25% each) and group 1 (14, 9.75%) contains predominantly species of the *Methylobacterium* genus.

**Figure 1.**
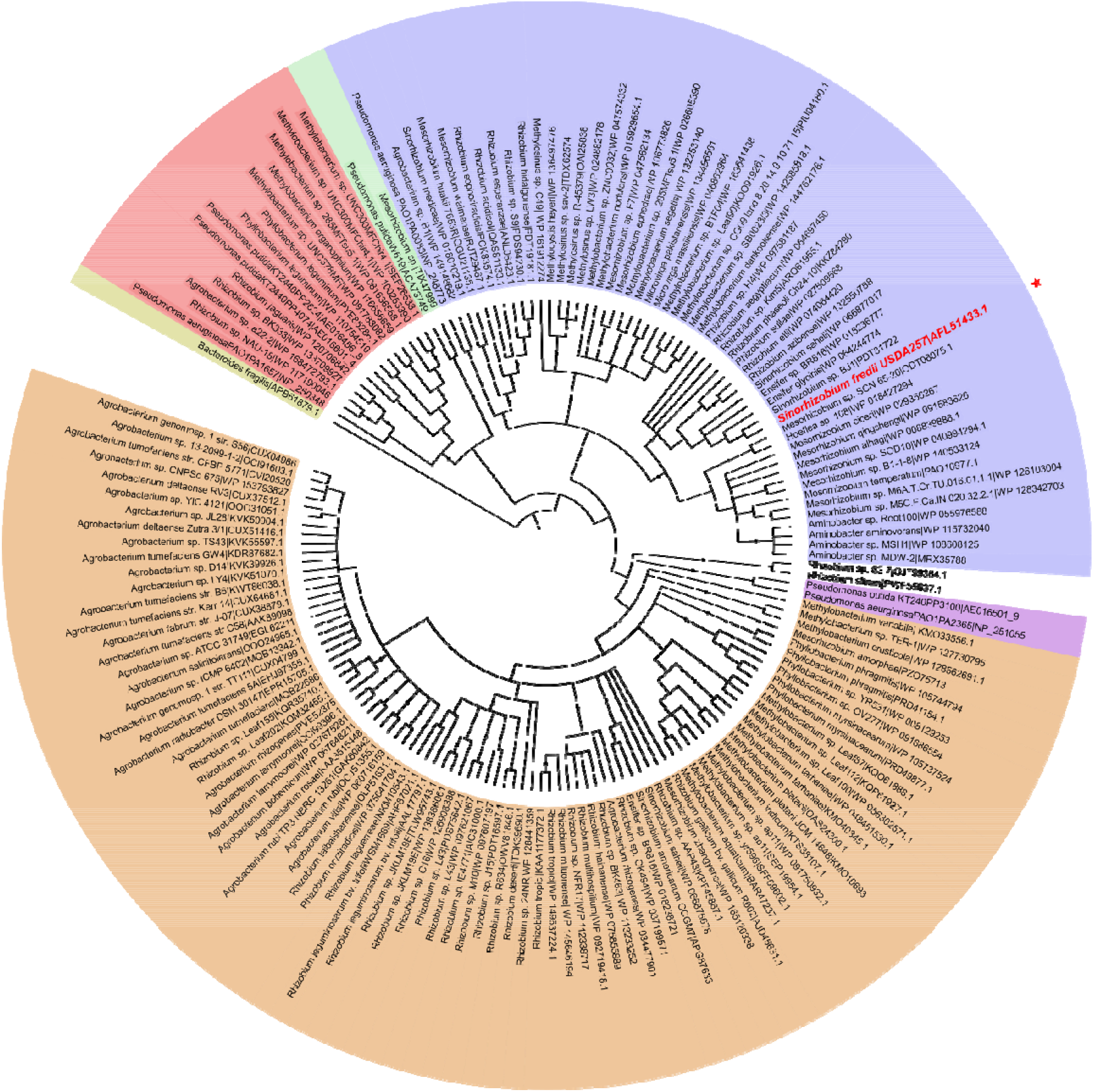
Phylogenetic tree of T6SSs of nitrogen-fixing bacteria. Maximum likelihood tree with 500 bootstrap replicates were built with Mega 7 software for the core component protein TssB. T6SS cluster nomenclature is used to show the main phylogenetic clusters (Boyer *et al*., 2009). Branches with black circles indicate a confidence level higher than 0.75. Five groups are distinguished: group 1 (red), group 2 (green), group 3 (blue), group 4 (purple) and group 5 (orange). Bacterium of the genus *Bacteroides* (brown) represents the tree root. Red star indicates the position of *S. fredii* USDA257 T6SS.

Some bacteria groups, namely *Pseudomonas,* frequently contain 2 or more T6SS clusters (Bernal *et al*, 2017). However, despite the number of T6SS clusters in a strain ranging from 1 to 5, in a broader group such a phytobacteria, only an estimate 7% of strains contain more than one cluster (Bernal *et al*., 2018). This number is even smaller among rhizobial T6SSs where most strains contain only a single T6SS cluster.

### 3.2. The reference rhizobial strain S. fredii USDA257 possesses a complete T6SS

Inspection of USDA257 genome revealed 26 T6SS-related ORFs located in a chromosomal cluster (**Fig. 2A**, **Table 1**). Thirteen genes (*tssABCDEFGHIJKLM*) encode the structural proteins required for a functional T6SS, including the membrane complex, the base plate, the tail and the ATPase that recycles the system. We further identified genes encoding a previously described regulatory phosphorylation cascade (Mougous *et al*., 2007), including the threonine kinase/phosphatase pair (PpkA-PppA) and the phosphorylation substrate (Fha). TagF is a posttranslational repressor that regulates T6SS via Fha interaction. TagF-linked T6SS repression have been studied in several systems, including *P. aeruginosa* and *A. tumefaciens* T6SSs (Lin *et al*., 2018). A putative TagF protein is encoded in USDA257 T6SS cluster, indicating that this system might have a similar posttranscriptional regulation to both *P. aeruginosa* and *A. tumefaciens*.

**Figure 2.**
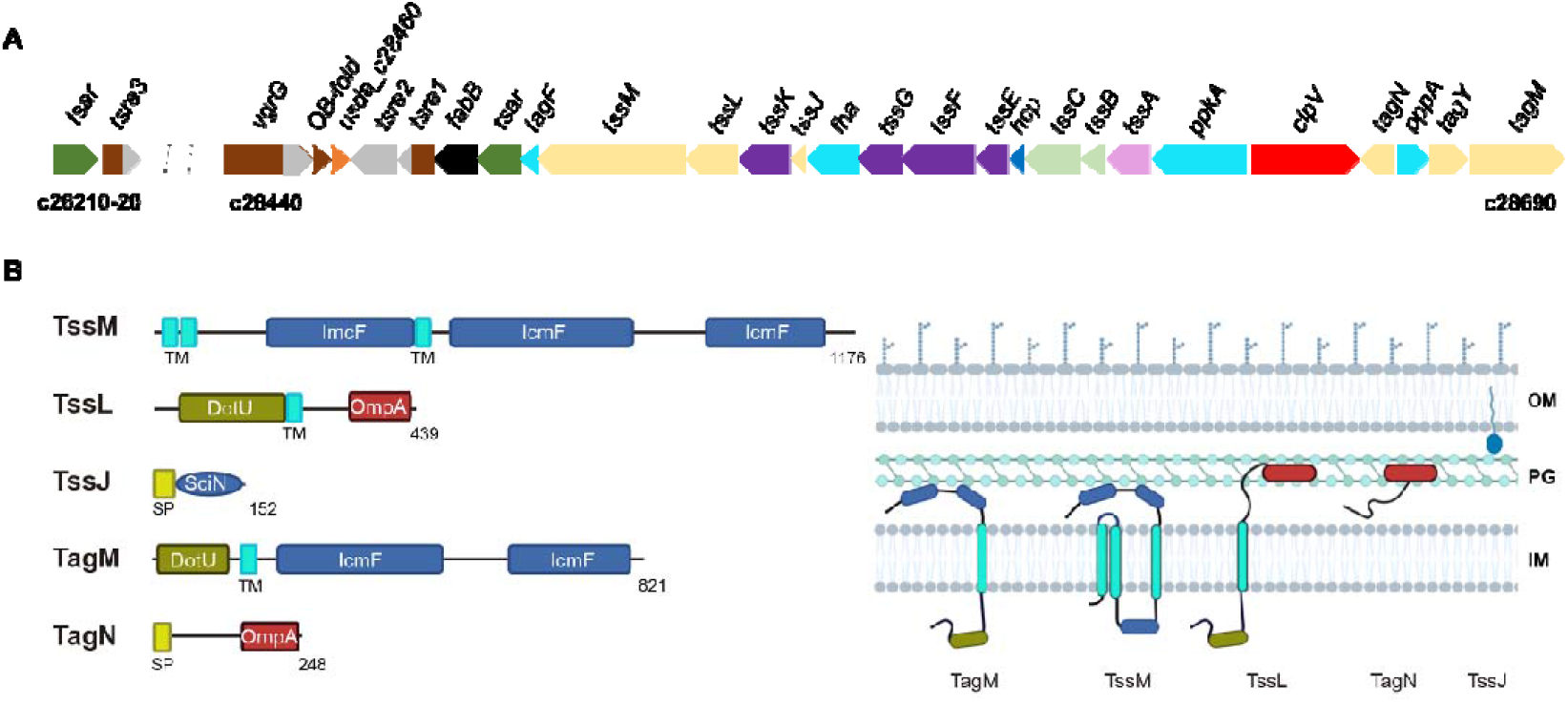
The T6SS of *S. fredii* USDA257. (A) Organization of the T6SS cluster of USDA257. Color codes: Genes that encode the components of membrane complex (*tssMLJ*) and accessory proteins (*tagLYM*) (veige). Genes coding for components of the base plate (*tssKGFE*) (purple). *tssA* (pink). Genes *tssB* and *tssC* (contractile sheath) (light green). The *hcp* gene, that encodes the protein forming the inner tube (dark blue). Genes that encode regulatory proteins (*tagF*, *fha*, *ppkA* and *pppA*) (light blue). *clpV*, that codes for ATPase (red). Genes coding for possible effectors of the system (C-term domain of *vgrG*, *tsre1, tsre2* and *tsre3*) (grey). Genes that encode a possible adapter (*tsar*) (dark green). Gene of unknown function (*usda_c28460*) (orange). The *vgrG* and *OB-fold* genes (brown). *paar* domain (brown). (B) Schematic representation of the structural protein domains of TssM, TssL, TssJ and the accessory proteins TagM and TagN. DotU cytosolic domain (dark green), transmembrane domain (light blue), IcmF structural domain (dark blue), OmpA peptidoglycan-binding domain (red), and signal peptide (light green). OM: outer membrane, IM: inner membrane, PG: peptidoglycan layer.

**Table 1.** Genes of the T6SS cluster of *S. fredii* USDA257.

| Locus name | Protein name | Conserved domains<sup>a</sup> | Celullar localization prediction |
| --- | --- | --- | --- |
| USDA257_c28440 | VgrGe | COG3501 (VgrG) + COG3210 (FhaB)/ pfam06715 (Gp5_C) | C |
| USDA257_c28450 | VgrG dom | COG3501 (VgrG)/ TIGR01646 (Vrg_GE) | D |
| USDA257_c28460 |  | TIGR02270 | C |
| USDA257_c28470 | Tre2 | Glycosyl hydrolase GH74 | C |
| USDA257_c28480 | Tre1 | PAAR-pfam13665 (DUF4150) | C |
| USDA257_c28490 | FabB | COG0304 (FabB) | MI |
| USDA257_c28500 | Tar | DUF2169 | C |
| USDA257_c28510 | TagF | COG3913 (SciT) | D |
| USDA257_c28520 | TssM | COG3523 (IcmF) | MI |
| USDA257_c28530 | TssL | COG3455 (DotU) + Pfam00691 (OmpA) | ME |
| USDA257_c28540 | TssK | COG3522 | C |
| USDA257_c28550 | TssJ | COG3521 | MI |
| USDA257_c28560 | Fha | COG3456 | C |
| USDA257_c28570 | TssG | COG3520 | C |
| USDA257_c28580 | TssF | COG3519 | C |
| USDA257_c28590 | TssE | COG3518 | C |
| USDA257_c28600 | Hcp | COG3157 (Hcp) | C |
| USDA257_c28610 | TssC | COG3517 | C |
| USDA257_c28620 | TssB | COG3516 | C |
| USDA257_c28630 | TssA | COG3515 + TIGR03363 | C |
| USDA257_c28640 | PpkA | COG0515 (SPS1) | P + PC |
| USDA257_c28650 | ClpV | COG0542 (ClpA) + pfam07724 (AAA_2) | C |
| USDA257_c28660 | TagL | COG2885 (OmpA) | ME |
| USDA257_c28670 | PppA | COG0631 (PTC) + TIGR02865 (spore_II_E) | C |
| **USDA257_c28680** | TagY | Pfam04102 (SlyX) +<br>pfam13539<br>(peptidasa_M15_4) | D |
| **USDA257_c28690** | TagM | COG3523 (IcmF) +<br>COG3455 (DotU) | D |

OM: outer membrane, IM: inner membrane, C: cytoplasmic, P: periplasm, PC: cell wall,
D: unknown, aa: amino acid.
a: conserved domains according to COG / pfam / TIGR / Phyre<sup>2</sup>.

TssJ, TssL and TssM are the core components of the T6SS membrane complex that docks the system to the cell envelope. TssL (DotU domain) and TssM (IcmF domains) are anchored to the inner membrane through one and three transmembrane helices respectively and form a channel through the cell envelope by interaction with each other and the outer membrane lipoprotein TssJ. Additional membrane-complex-related proteins have been described in some systems, including a set of proteins that anchor the T6SS to the peptidoglycan (PG) through the PG-binding domain OmpA (pfam00691) (Aschtgen *et al*., 2010). The set included the specialized TssL with a canonical N-terminal DotU domain and a C-terminal OmpA domain that have been identified in *P. aeruginosa* and *A. tumefaciens* among others. The authors also described a ∼250 amino acids protein with an N-terminal signal peptide and a C-terminal OmpA domain identified in *Burkholderia pseudomallei* and *Ralstonia solanacearum* and named TagN. The localization, topology, PG-binding, and interactions with the rest of the membrane complex remains to be characterized for this T6SS protein.

An in-depth analysis of USDA257 T6SS components, we identified the three core components of the membrane complex *i.e.,* TssJ, TssL and TssM, and two related accessory proteins named, TagM and TagN (**Fig. 2B**). In addition, USDA257 TssJ and TssM proteins contain the conventional T6SS domains, while USDA257 TssL is a specialized TssL (TssL_e_). We identified two additional accessory components, a TagN homologue and TagM, an 821 amino acids protein that shares certain degree of homology with both TssM and TssL (**Fig. S2**). This protein contains an N-terminal DotU domain, a single transmembrane domain and two IcmF domains; its transmembrane domain and the absence of a specific signal peptide indicates that this protein might be anchored to the inner membrane. USDA257 membrane complex varies from the canonical, presenting not only one but two membrane complex proteins with PG-binding domains (TssL_e_ and TagN) and a TssM-TssL hybrid protein (TagM) (**Fig. 2B**). This complexity might indicate a different assembling mode/capacity to anchor to the cell envelope.

Lastly, we have identified a gene encoding an additional accessory protein we have named *tagY* (**Fig. 2A**). TagY presents homology with an M15 peptidase, a D-alanyl D- alanine carboxypeptidase that is involved in bacterial cell wall biosynthesis (Lessard *et al*., 1999). Predictions indicate that TagY has a transmembrane domain at the N- terminal end and depending on the software used, it might have a signal peptide, suggesting that this protein could be anchored to the outer membrane with the C- terminal end facing the periplasm or the cellular exterior.

### 3.3. The S. fredii USDA257 T6SS cluster harbors genes encoding potential effectors

Frequently, T6SS effectors are genetically linked to *vgrG* and *hcp* genes, which can be found within a T6SS cluster or, in different numbers, scattered through the genome. For instance, *P. aeruginosa* PAO1 genome contains 10 *v*gr*G* and 5 *hcp* genes (Hachani *et al*., 2016). The *in silico* study of the USDA257 genome has revealed single copies of *vrgG* and *hcp* genes in the T6SS cluster (**Fig. 2A**). *hcp* is found surrounded by the structural genes *tssC* and *tssE* and there is no evidence of genes encoding putative effectors in the *hcp* proximity. On the other hand, *vrgG* is located at the end of the cluster and genetically linked to putative T6SS effectors and adaptors, displaying a similar genetic architecture to *P. aeruginosa* PAO1 and *P. fluorescens* F113 *vrgG1b* clusters (Hachani *et al*., 2011; Redondo-Nieto *et al*., 2013). These clusters encoded an evolved VrgG, an oligonucleotide-binding(OB)-fold, a DUF2169 adaptor (an immunoglobulin-like protein), a thiolase-like protein (PRK06147) proposed to be a novel T6SS adaptor, a PAAR protein with a C-terminal cytotoxic domain, an immunity protein and a heat repeat-containing protein. Both sets of genes only differ in the sequence of the toxic domain and the immunity pair, a common characteristic of this genetic island, previously described by Pissaridaou *et al*. (2018). In USDA257, genes downstream *vrgG* are inverted, but aside from that, the genetic organization of the region is conserved compared to previously described ones. Thus, USDA257 contains the genes encoding an evolved VrgG with a similar C-terminal domain, the OB-fold protein, the DU2169 adaptor named *tsar* (type <u>s</u>ix <u>a</u>dapter rhizobium), followed by a thiolase-like protein (T6SS adaptor), an evolved PAAR protein named Tsre1 (*tsre*, type <u>s</u>ix rhizobial <u>e</u>ffector) with a C-terminal domain of unknown function and, lastly, a protein named Tsre2 (**Fig. 2A**, **Tables 1 and 2**). According to Phyre^2^, in which model is built based on the experimental structure of a homologue that serves as a structural template (Kelley *et al*., 2015), Tsre2 shows structural homology with a glycoside hydrolase enzyme GH74 with a xyloglucan binding domain from *Caldicellulosiruptor lactoaceticus* 6A (69% alignment, 99.3% confidence). In addittion, the predicted model of the USDA257 Tsre2 protein was collected from the Alphafold database and queried using the FoldSeek tool, that predict the structural homology based on the terciary interactions of proteins in a sequence-independent manner (Van Kempen *et al*., 2023). Using this approach, best prediction for the structural homology is found with the immune protein Tsi7 from *P. aeruginosa* PAO (probable structural homology between amino acid residues 2 and 340 with an E-Value of 3.77e-14), whereas the structural homology with the glycoside hydrolase enzyme ranges between residues 11 and 338 (1.10e-1 E-Value). Interestingly, the evolved PAAR protein named Tsre1, which does not display any structural homology in its C-terminal domain according to Phyre^2^, is predicted to be homologous to the DNase toxin Tse7 of *P. aeruginosa* PAO1 (ranging from 3 to 321 amino acid positions, 5.25e-24 E-Value).

**Table 2.** Best predictions for the structural homology from representative rhizobial versions coded by same loci than *tsre1* and *tsre2* genes of *S. fredii* USDA257.

| **Rhizobial strain** | **Tsre1-like** | **Tsre2-like** |
| --- | --- | --- |
| *Sinorhizobium fredii* USDA257 | P: PAAR (25-125 a.a.)<br>F: DNase toxin Tse7 (3-321 a.a.)<br>Size: 324 a.a. | P: Glycosyl hydrolase GH74 (1-340 a.a.)<br>F: Immune protein Tsi7 (2-340 a.a.)<br>Size: 351 a.a. |
| *Rhizobium phaseoli* N841 | P: PAAR (26-126 a.a.).<br>F: DNase toxin Tse7 (1-327 a.a.)<br>Size: 328 a.a. | P: Glycosyl hydrolase GH74 (106-301 a.a.)<br>F: Immune protein Tsi7 (17-228 a.a.)<br>Size: 309 a.a. |
| *Rhizobium phaseoli* R620 | P: PAAR (26-126 a.a.).<br>F: Not available.<br>Size: 330 a.a. | P: Glycosyl hydrolase GH74 (98-302 a.a.)<br>F: Not available.<br>Size: 309 a.a. |

Structural homologies were determined by Phyre<sup>2</sup> (P:) software and (F:) Foldseek tool.
In parenthesis are indicated range of aligned amino acids.

Not only the *paar* genes of USDA257 but also *vgrG* gene encode for evolved proteins, in which C-terminal ends contain putative effector domains (Pukatzki *et al*., 2009). According to sequence similarity, the C-terminal domain of VgrG corresponds to a 225 amino acid fragment from the FhaB protein, a chaperone that prevents the premature folding of hemagglutinin filaments, one of the virulence factors of *Bordetella pertussis* (Coutte *et al*., 2001). However, the analysis performed with Phyre^2^ software indicates that this C-terminal domain also shows structural homology with the Cthe_2159 protein from *Hungateiclostridium thermocellum* ATCC 27405 (85% alignment, 86.5% confidence), a polysaccharide lyase-type protein that binds to cellulose and polygalacturonic acid (**Fig. 2A**, **Table 2**). The predicted model of the evolved VrgG from USDA257 using Foldseek determined that the *Nitrospirae bacterium* structural homologous VgrG is the most similar protein among all proteins (between amino acid residues 3 and 768, 1.40e-75 E-Value).

Interestingly, we identified one potential orphan effector, since Tsar adaptor and evolved PAAR proteins, termed Tsre3, were found separately from the T6SS cluster (**Fig. 2A**). Curiously, this orphan PAAR protein displays sequence dissimilarity with the Tsre1 effector both in the N-terminal and C-terminal domains and does not exhibit structural homology with any previously described proteins according to Phyre^2^. Using FoldSeek, the structure of a DUF4150 domain-containing protein with unknown function from *Methilobacterium* sp. yr596 is identified with probable structural homology only with the N-terminal part (amino acid residues from 38 to 168 with a E- Value 1.11e-16).

### 3.4. The vrg cluster is well conserved among rhizobia except for the effector region

To obtain a broader perspective of functioning and distribution of *tsre1* and *tsre2* genes among rhizobia, we performed a comparison of the *vgrG* clusters from 18 rhizobial strains. **Fig. 3** shows sequence conservation across the entire cluster. As expected, the genes encoding the VgrG and the OB-fold proteins are conserved and the region encoding the putative proteins Tsre1 and Tsre2 is the one varying the most. Curiously, the 3′end of the gene encoding the DUF2169 adaptor and 5′end the gene encoding the thiolase-like protein are less conserved and could indicate that they are effector/toxin- specific.

**Figure 3.**
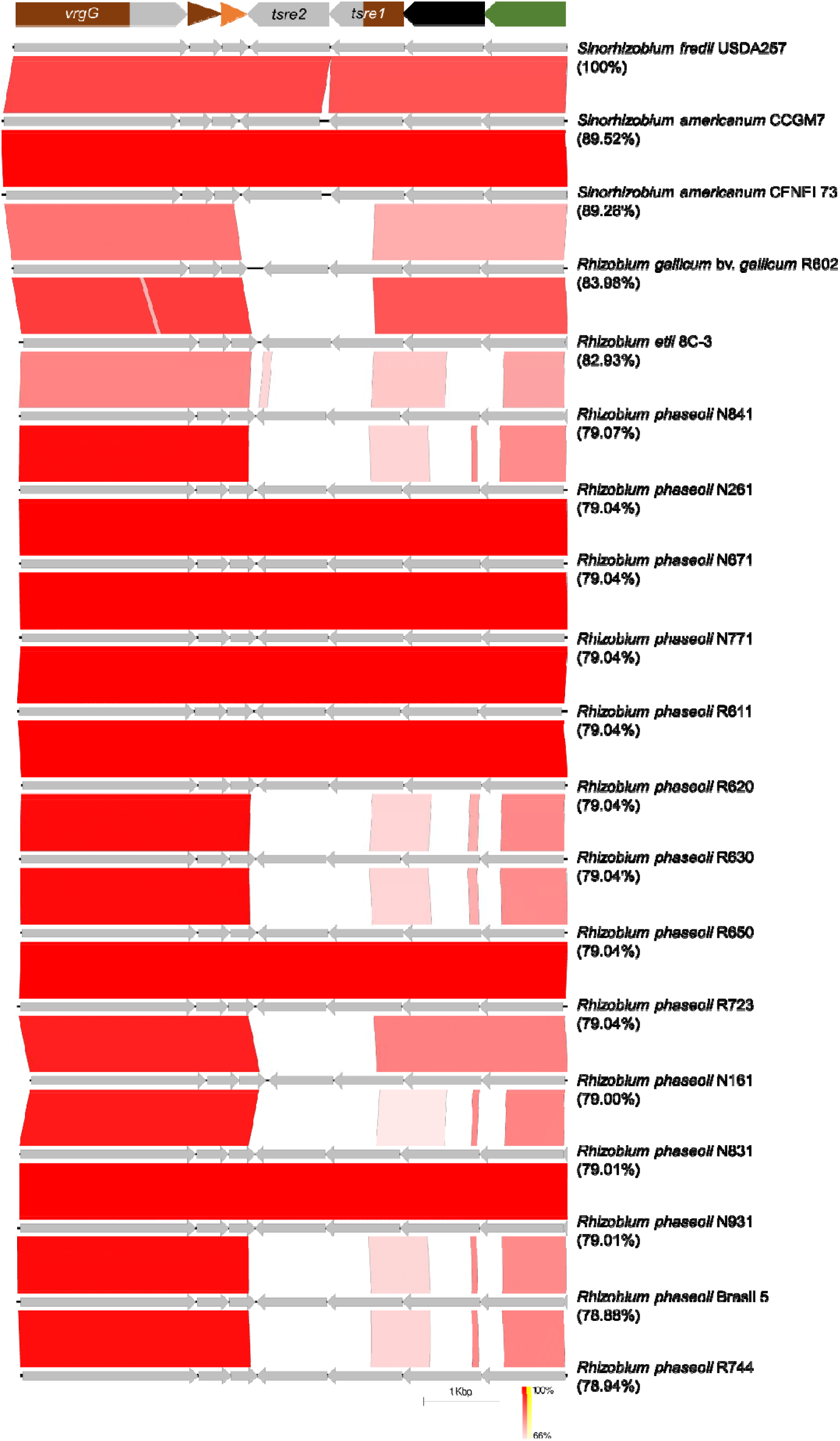
Genome sequence alignment of the *vgrG* region demonstrating the divergence of *tsre1* and *tsre2* in eighteen rhizobial strains using Easyfig. Clusters are sorted by homology degree respect to the VgrG protein from *S. fredii* USDA257 (Blastp E value). In parenthesis is indicated the percentage of identity for each VrgG. Putative protein domains encoded for each gene of the *vgrG* region are boxed above alignments.

A neighbor-joining phylogeny of the Tsre1 and Tsre2 proteins from these rhizobial strains clearly group USDA257 with *Sinorhizobium americanum* strains CCGM7 and CFNEI 73 for both proteins, whereas most Tsre1 and Tsre2 *Rhizobium phaseoli* versions are distributed in two separated branches, one grouping versions from strains R620, R650, R611, N771, N671, N261, R723 and Brasil 5, and the other one clustering proteins from strains N841, R630, N831, N931 and R744 (**Fig. 4**). Alignment of the Tsre1 protein sequences indicated a high degree of similarity among the N-terminal but not in the C-terminal domains in all versions, that were grouped as in the phylogenetic tree (**Fig. S3-5**). In this case, the first half of the three versions of Tsre1 proteins showed a clear structural homology with PAAR domain according to Phyre^2^ (**Table 2**). Despite alignment of Tsre2 versions showeing a lower degree of similarity among proteins from different branches of the phylogenetic tree (**Fig. S6-8**), the structural homology predicted using this approach indicates that the other two versions are structurally similar than the same GH74 glycoside hydrolase enzyme, differencing in their range of alignment. When using the Foldseek tool, the proteins most structurally similar to the N841 versions of both Tsre1 and Tsre2 are Tse7 and Tsi7 from *Pseudomonas aeruginosa* PAO1. In contrast, for the R620 version, the lack of an AlphaFold prediction made structural comparison of the protein impossible using this approach (**Table 2**).

**Figure 4.**
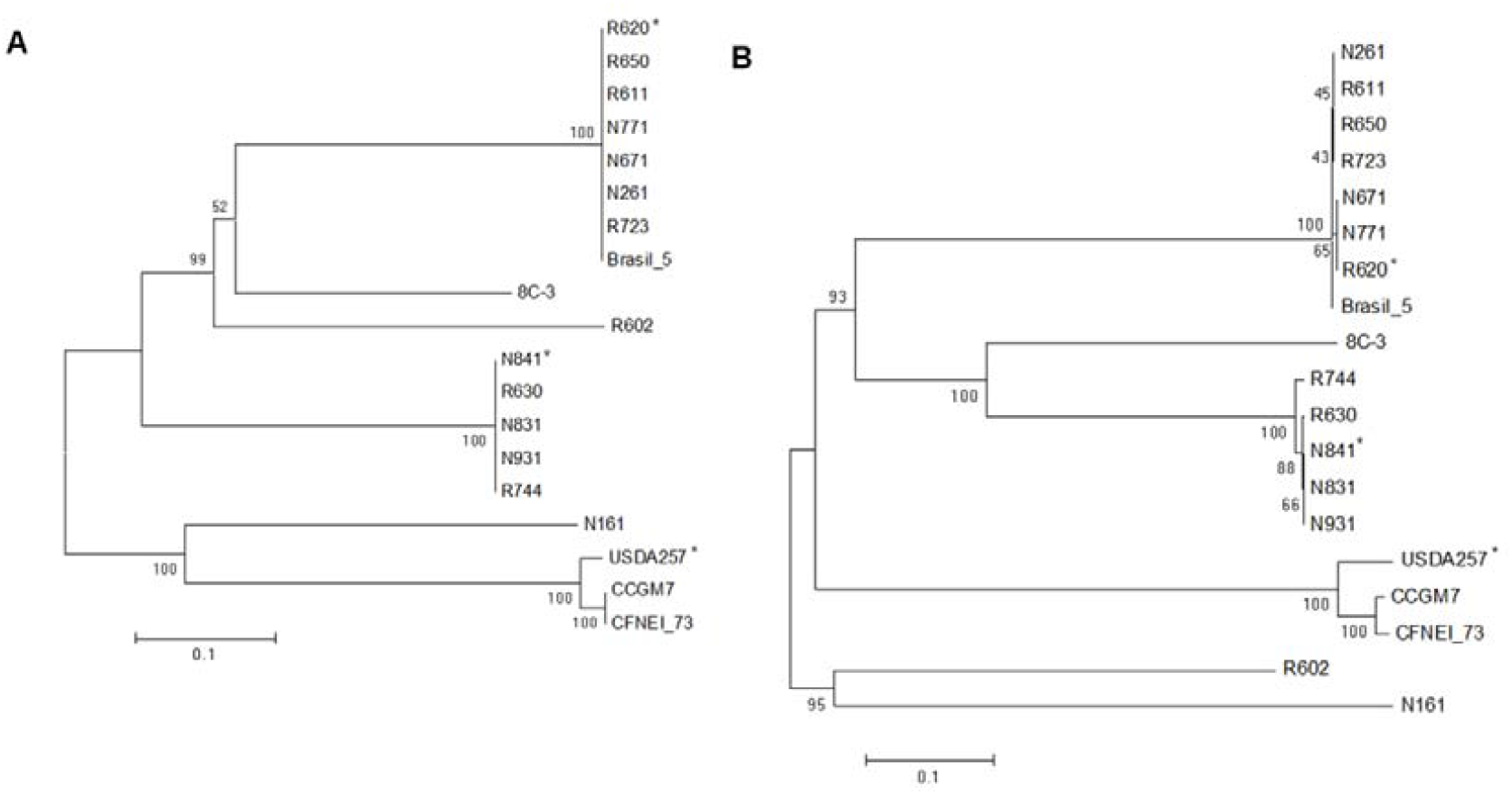
Neighbor-joining tree of rhizobia T6SS putative effectors. Bootstrap values ≥40 is indicated for each node. The cluster analysis to group the strains by Tsre1 (A) or Tsre2 (B) sequence similarities was done using the program CLUSTALW in the MEGA7 software package with the algorithm neighbor-joining method. Tree robustness was assessed by bootstrap resampling (1000 replicates each). Asterisks indicate representative Tsre1 and Tsre2 versions for each main branch.

### 3.5. The T6SS of S. fredii USDA257 is functional and induced in minimal medium and in the nodules

To determine the conditions under which the T6SS of *S. fredii* USDA257 is expressed, we cloned the promoter region of the first gene of the T6SS structural operon (*ppkA*) upstream of the reporter gene *lacZ*. Thus, by ß-galactosidase assays, we aimed to measure the expression levels of the USDA257 T6SS genes. We tested USDA257 cultures 30 hours post inoculation in selected rich, standard and minimal media for growing rhizobia: TY, YM, and MM respectively (**Fig. 5A**). YM is a standard medium possessing moderate quantities of yeast extract that supply the limiting organic nitrogen and micronutrients, whereas MM is a minimal medium that only has glutamate as nitrogen source. Both YM and MM media can be prepared with different concentrations of mannitol as carbon source. Thus, the activation pattern in the ß-galactosidase assays of the wild-type strain carrying the construct containing the USDA257 *ppkA* promoter region fused to the *lacZ* reporter gene indicated that the expression of the T6SS genes is highest in the minimal medium MM3 (3 g L^-1^ of mannitol), whereas the lowest values were obtained when this rhizobium was grown in the rich medium TY (about 3-fold and 1,8-fold, respectively, in comparison to those values obtained by the strain carrying the empty plasmid) (**Fig. 5A**). To confirm these findings, we followed a complementary approach, measuring the transcriptional activation of the USDA257 *ppkA* gene. *q*RT- PCR experiments shed the same transcriptional activation trend for the *ppkA* gene in USDA257 cultures after 48 hours post inoculation. The highest transcriptional values were found in cultures growing at MM3 medium (around 8-fold) in comparison to the values obtained for the same genes in TY medium (**Fig. 5B**). To delve further into the regulation of T6SS, we performed ß-galactosidase assays at different points of the USDA257 growth curve, inoculating the bacterium in the MM3 inducing-medium.

**Figure 5.**
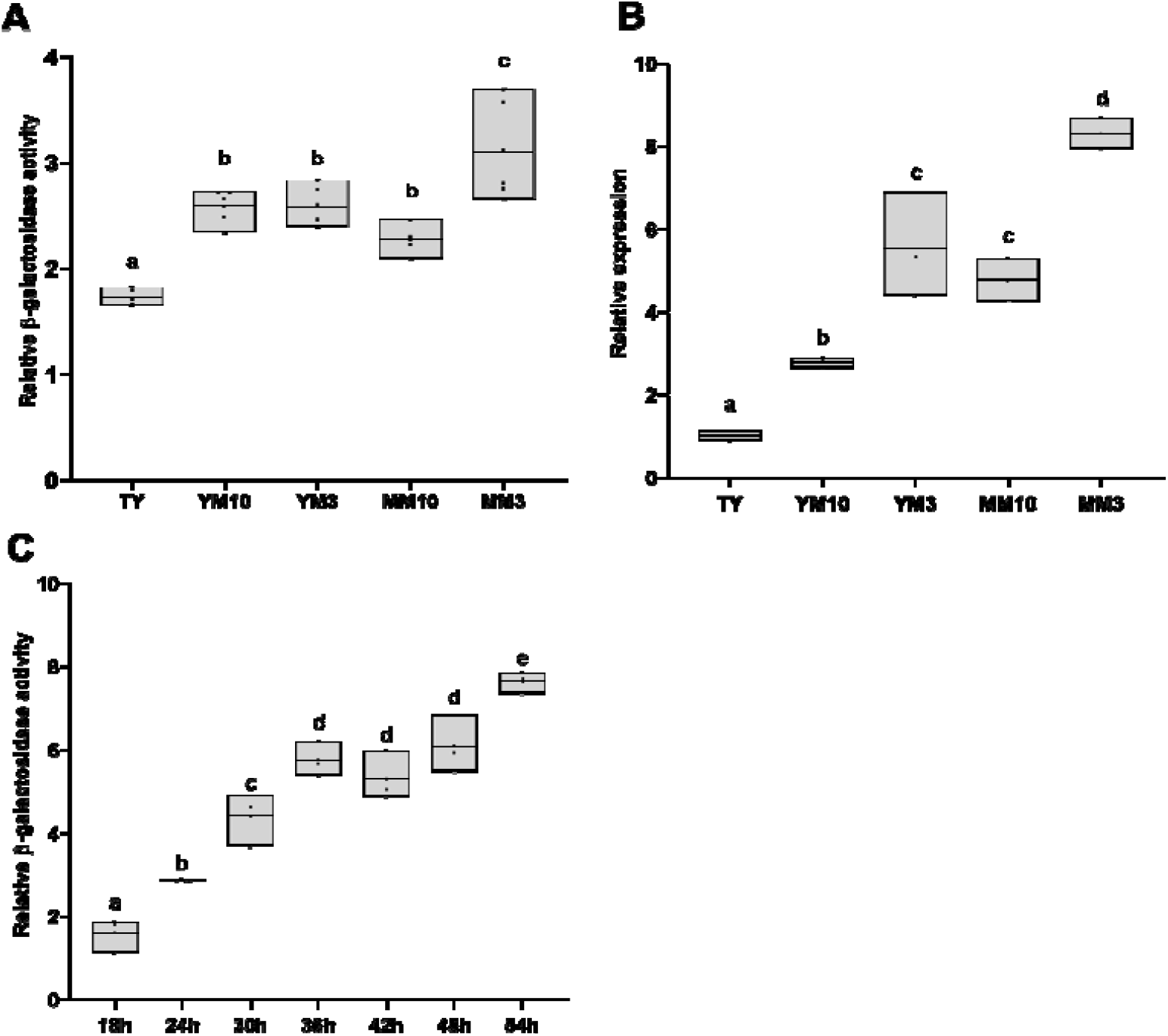
Expression of the *S. fredii* USDA257 *ppkA* gene. (A) Fold-change values of the β-galactosidase activity of USDA257 carrying a pMP220 plasmid containing the *ppkA* promoter region fused to the *lacZ* gene respect to those values obtained by the strain carrying the empty pMP220 plasmid. Assayed conditions were TY, YM10, YM3, MM10 and MM3 media 30 hours after inoculation. Data followed by the same letter are not significantly different at the level of α = 5% (One-Way ANOVA with multiple comparisons, P<0.05). (B) Quantitative RT-PCR analysis of the expression of the *ppkA* gene of USDA257 grown in the TY, YM10, YM3, MM10 and MM3 media 48 hours after inoculation. Expression data shown are the mean (± standard deviation of the mean) for three biological replicates. Data followed by the same letter are not significantly different at the level of α = 5% (One-Way ANOVA with multiple comparisons, P<0.05). (C) Fold-change values of the β-galactosidase activity of USDA257 carrying a pMP220 plasmid containing the *ppkA* promoter region fused to the *lacZ* gene respect to those values obtained by the strain carrying the empty pMP220 plasmid. The assayed condition was MM3 medium at different hours after inoculation. Data followed by the same letter are not significantly different at the level of α = 5% (One-Way ANOVA with multiple comparisons, P<0.05).

Interestingly, the activation pattern of the wild-type strain carrying the P_ppkA_::*lacZ* fusion pointed out that the expression of the T6SS follows the archetypical growth curve profile, with a saturation of the ß-galactosidase values around 54 hours after inoculation (about 7-fold in comparison to those values obtained by the strain carrying the empty plasmid) (**Fig. 5C**).

To gain insight about the potential role of the USDA257 T6SS in the early steps of the symbiotic process, β-galactosidase assays were performed in the presence of genistein, a *nod*-gene inducing flavonoid in several related *S. fredii* strains (Jiménez-Guerrero *et al*., 2015, 2018; Pérez-Montaño *et al*., 2014). Activation of the *ppkA* promoter was measured in MM3 medium at stationary phase in the presence or the absence of genistein, using as control of *nod*-gene induction the wild-type strain carrying the pMP240 plasmid, which harbors the conserved *nodA* promoter of *R. leguminosarum* bv. *viciae* fused to the *lacZ* reporter gene. In the presence of the USDA257 *nod* gene- inducing flavonoid genistein, induction values of strains carrying the *ppkA* promotor fused to *lacZ* were similar among conditions (**Fig. S9**), indicating that the expression of the T6SS is not controlled by the presence of this symbiotic-plant molecule.

The extracellular secretion of the Hcp protein into a culture medium is a hallmark to assess functionality of the T6SS system. Therefore, we produced a specific polyclonal antibody against USDA257 Hcp to assess USDA257 T6SS activity in the previously described conditions, *i.e.* the different culture media after 48 hours of growth (**Fig. 6A**). As anticipated, Western blot analysis corroborated the results obtained from both transcriptomic assays, revealing that the presence of Hcp was lowest in TY medium and highest in both MM media. We also used a USDA257 *tssA* mutant strain to disable the system, since TssA is an essential component for T6SS activity. As expected, we detected Hcp in the supernatant of the wild-type strain grown in MM3 but not in the isogenic *tssA* mutant, while it was present in the cytosol of both strains (**Fig. 6B**). Altogether, results stablishe that USDA257 T6SS is a functional secretion machinery, and it is found most active in poor media at the stationary phase of growth.

**Figure 6.**
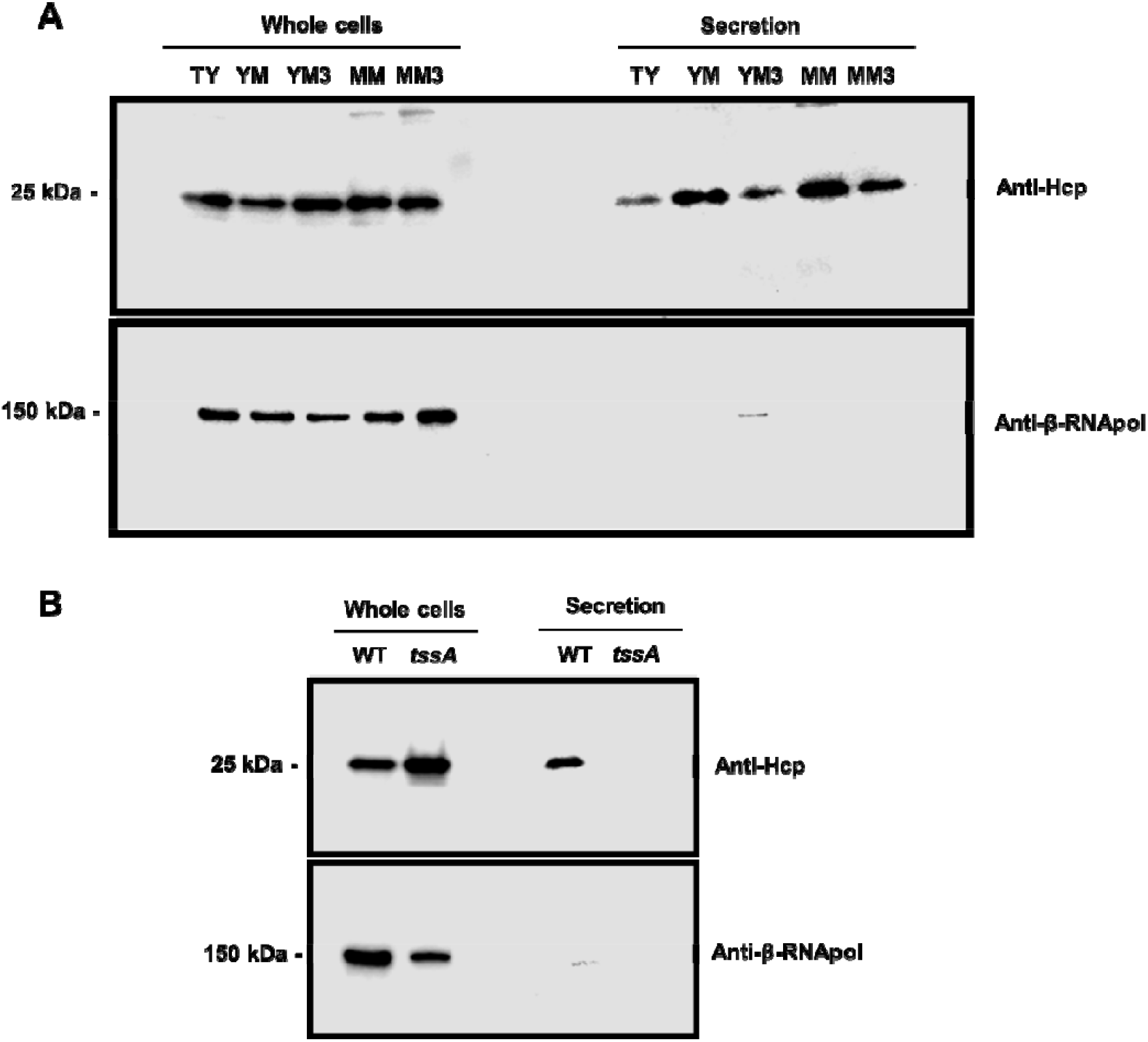
Functionality of the *S. fredii* USDA257 T6SS. (A) Production and secretion of Hcp in the USDA257 wild-type strain grown for 48 h in rich, standard and minimal media: TY, YM, and MM, repectively. YM3 and MM3 represent derivatives of their corresponding media with reducedi carbon source content,. (B) Production and secretion of Hcp in the USDA257 wild-type and the *tssA* mutant strains grown in MM3. For both panels, the Hcp protein was detected by Western blot analysis using a specific anti-Hcp antibody. Detection of the β-subunit of the RNA polymerase (β-RNApol) was used as control. “Whole cells” represents the intracellular protein fraction, whlilThe position of the molecular size marker (in kDa) is indicated.

Finally, to elucidate the USDA257 T6SS potential functioning specifically during nodule colonization in the rhizobial symbiosis, we conducted a detailed investigation by confocal microscopy into the localization of rhizobial cells within nodules, along with the activation pattern of their T6SS. The expression of both constitutive GFP and T6SS- responsive RFP allowed to account for all dual-bioreporter cells *in vivo* during the infection of legume nodules. Interestingly, strong constitutive GFP and T6SS- responsive RFP fluorescence expression were observed for all rhizobial cells that are colonizing the symbiotic cells of *L. burttii* nodules (**Fig. 7C**), which reveales that the T6SS of USDA257 is fully active when rhizobial cell invade the host nodule tissue.

**Figure 7.**
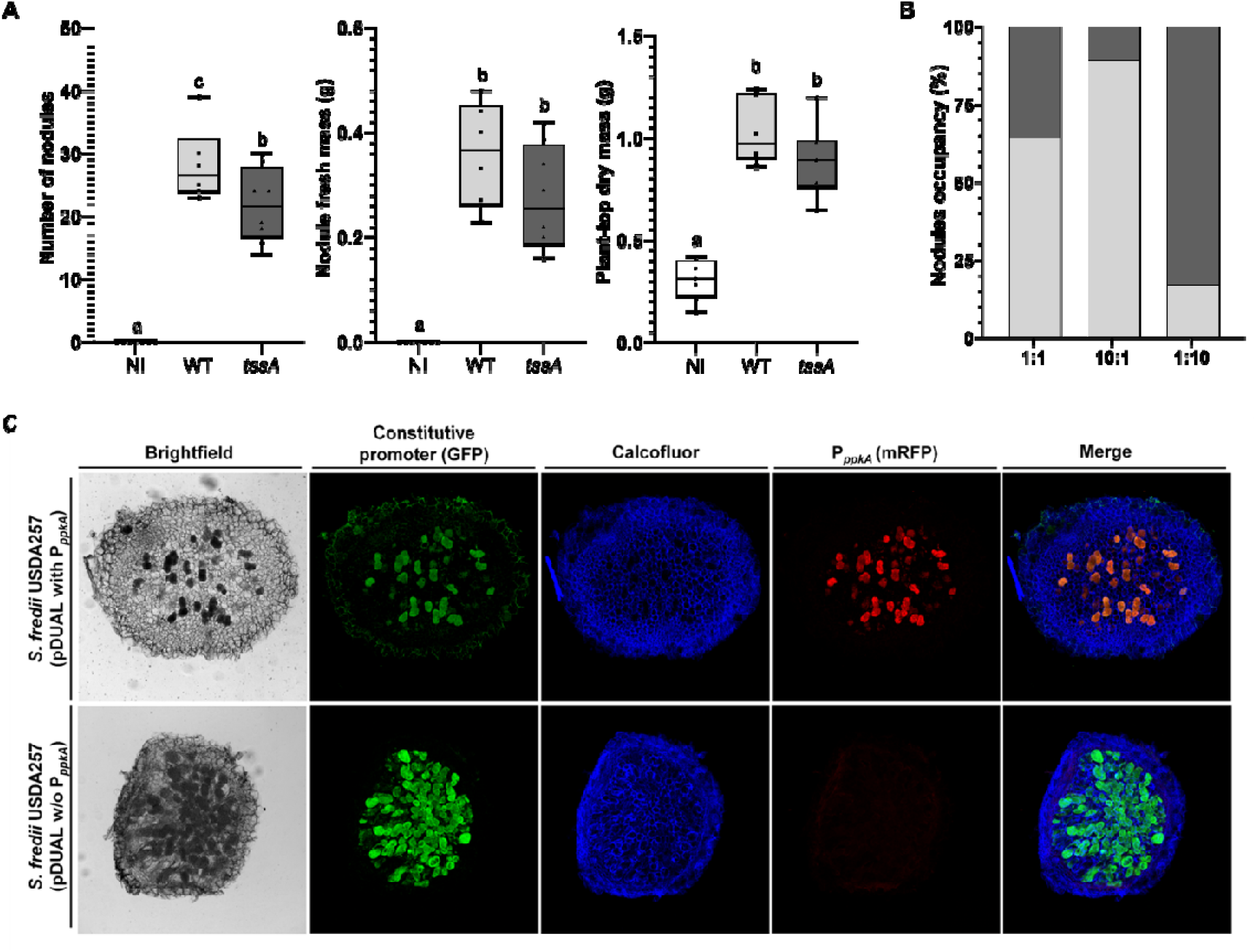
Plant responses to inoculation with *S. fredii* USDA257 strains. (A) Plant responses to inoculation of *G. max* cv Pekin with different *S. fredii* USDA257 strains. Plants were evaluated 6 weeks after inoculation. NI: non-inoculated plants. Data followed by the same letter are not significantly different at the level of α = 5% (One-Way ANOVA with multiple comparisons, P<0.05). (B) Competition for nodulation assays of different *S. fredii* USDA257 strains. Percentages of nodule occupancy were evaluated by co-inoculation at different ratios of both strains at 30 dpi. Light grey: USDA257 wild-type strain, dark grey: USDA257 *tssA* mutant strain. (C) Confocal microscopy of *L. burttii* 30 dai-nodules infected by the *S. fredii* USDA257 carrying a dual fluorescent reporter plasmid with (pDUAL with P_ppkA_) or without (pDUAL w/o P_ppkA_) the *ppkA* promoter region upstream of the *rfp* gene.

### 3.6. The T6SS of S. fredii USDA257 is required for a successful nodulation and competitivenes with G. max cv Pekin

Due to many bacterial T6SS are mainly involved in interbacterial competition, we first performed interbacterial competition assays. These experiments we carried out between the wild-type and a *ttsA* mutant (T6SS-defective) strains of USDA257 as predators and other bacterial species (*A. tumefaciens*, *P. carotovorum* and *S. fredii* HH103) as preys. The bacteria were mixed 1:1 (predator:prey) and co-cultured on USDA257 T6SS- inducing medium for 24 and 48 hours. Spots containing the competing bacteria were collected and the bacterial CFUs were determined on selective plates. The number of predator and prey bacteria after competing is shown in **Table 3**. In addition to the fact that no differences were observed between the wild-type and mutant strains in all conditions, both wild-type and the *tssA* mutant strains displayed a reduced number of bacteria when competing with any prey. These results may indicate that the USDA257 T6SS is not involved in interbacterial competition *in vitro* against another potential rhizospheric competitor as the pathogenic *A. tumefaciens* and *P. carotovorum* or the symbiont *S. fredii* HH103 (Bernal *et al*., 2017).

**Table 3.**
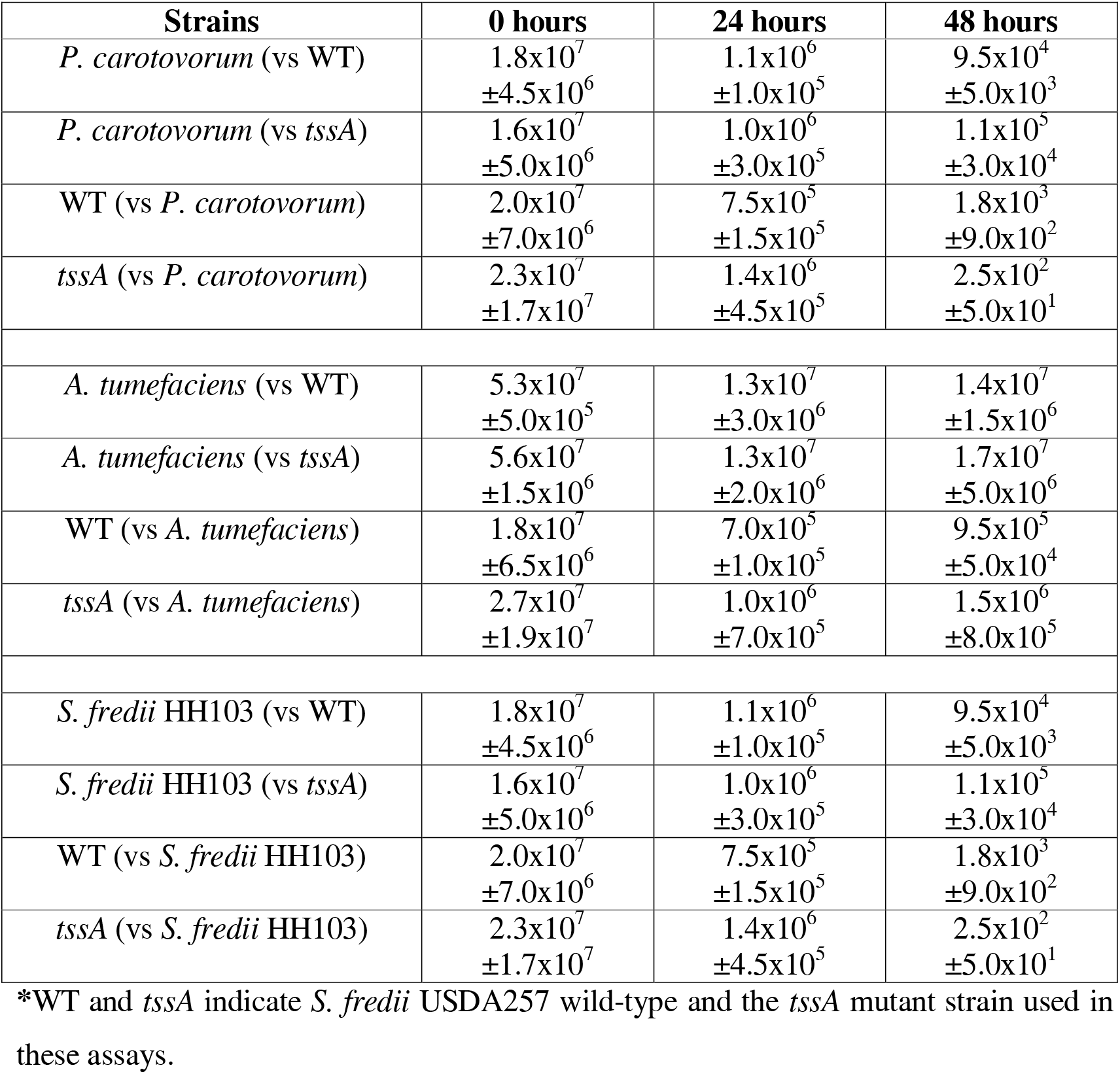
Average of CFUs obtained in competition assays after 0, 24 or 48 hours of interaction in T6SS-inducing media.

| **Strains** | **0 hours** | **24 hours** | **48 hours** |
| --- | --- | --- | --- |
| *P. carotovorum* (vs WT) | $1.8 \times 10^7$<br>$\pm 4.5 \times 10^6$ | $1.1 \times 10^6$<br>$\pm 1.0 \times 10^5$ | $9.5 \times 10^4$<br>$\pm 5.0 \times 10^3$ |
| *P. carotovorum* (vs *tssA*) | $1.6 \times 10^7$<br>$\pm 5.0 \times 10^6$ | $1.0 \times 10^6$<br>$\pm 3.0 \times 10^5$ | $1.1 \times 10^5$<br>$\pm 3.0 \times 10^4$ |
| WT (vs *P. carotovorum*) | $2.0 \times 10^7$<br>$\pm 7.0 \times 10^6$ | $7.5 \times 10^5$<br>$\pm 1.5 \times 10^5$ | $1.8 \times 10^3$<br>$\pm 9.0 \times 10^2$ |
| *tssA* (vs *P. carotovorum*) | $2.3 \times 10^7$<br>$\pm 1.7 \times 10^7$ | $1.4 \times 10^6$<br>$\pm 4.5 \times 10^5$ | $2.5 \times 10^2$<br>$\pm 5.0 \times 10^1$ |
| *A. tumefaciens* (vs WT) | $5.3 \times 10^7$<br>$\pm 5.0 \times 10^5$ | $1.3 \times 10^7$<br>$\pm 3.0 \times 10^6$ | $1.4 \times 10^7$<br>$\pm 1.5 \times 10^6$ |
| *A. tumefaciens* (vs *tssA*) | $5.6 \times 10^7$<br>$\pm 1.5 \times 10^6$ | $1.3 \times 10^7$<br>$\pm 2.0 \times 10^6$ | $1.7 \times 10^7$<br>$\pm 5.0 \times 10^6$ |
| WT (vs *A. tumefaciens*) | $1.8 \times 10^7$<br>$\pm 6.5 \times 10^6$ | $7.0 \times 10^5$<br>$\pm 1.0 \times 10^5$ | $9.5 \times 10^5$<br>$\pm 5.0 \times 10^4$ |
| *tssA* (vs *A. tumefaciens*) | $2.7 \times 10^7$<br>$\pm 1.9 \times 10^7$ | $1.0 \times 10^6$<br>$\pm 7.0 \times 10^5$ | $1.5 \times 10^6$<br>$\pm 8.0 \times 10^5$ |
| *S. fredii* HH103 (vs WT) | $1.8 \times 10^7$<br>$\pm 4.5 \times 10^6$ | $1.1 \times 10^6$<br>$\pm 1.0 \times 10^5$ | $9.5 \times 10^4$<br>$\pm 5.0 \times 10^3$ |
| *S. fredii* HH103 (vs *tssA*) | $1.6 \times 10^7$<br>$\pm 5.0 \times 10^6$ | $1.0 \times 10^6$<br>$\pm 3.0 \times 10^5$ | $1.1 \times 10^5$<br>$\pm 3.0 \times 10^4$ |
| WT (vs *S. fredii* HH103) | $2.0 \times 10^7$<br>$\pm 7.0 \times 10^6$ | $7.5 \times 10^5$<br>$\pm 1.5 \times 10^5$ | $1.8 \times 10^3$<br>$\pm 9.0 \times 10^2$ |
| *tssA* (vs *S. fredii* HH103) | $2.3 \times 10^7$<br>$\pm 1.7 \times 10^7$ | $1.4 \times 10^6$<br>$\pm 4.5 \times 10^5$ | $2.5 \times 10^2$<br>$\pm 5.0 \times 10^1$ |

\*WT and *tssA* indicate *S. fredii* USDA257 wild-type and the *tssA* mutant strain used in these assays.

To gain insight into the role and functioning of the *S. fredii* USDA257 T6SS, the symbiotic abilities of the wild-type or its derivate *tssA* mutant strains with *G. max* cv Pekin, considered its natural host, were inspected (**Fig. 7A**). Interestingly, the number of nodules in examined plants inoculated by the *tssA* mutant was statistically lower than those obtained in plants infected by the wild-type strain. Similar but not statistically trends were observed for both nodule fresh mass and plant-top dry mass parameters. On the other hand,the impact of the T6SS on the bacterial competitive nodulation capacity with *G. max* cv Pekin was assessed by co-inoculation experiments with USDA257 and the *tssA* mutant. In all ratios (1:1, 1:10 and 10:1), the T6SS defected strain was clearly outcompeted by the wild-type (**Fig. 7B**). Thus, results obtained from nodulation assays and competitiveness for nodule occupancy indicate that the T6SS of USDA257 is required for an optimal nodule development in soybean.

## 4. Discussion

Application of symbiotic rhizobia is a cheap and ecofriendly alternative source of nitrogen for plants, which favors a moderate use of nitrogen fertilizers mitigating their excessive utilization (Martinelli *et al*., 2020; Pérez-Montaño *et al*., 2014). The efficiency of symbiosis depends on various factors, such as the soil, the environment, and the symbiotic pair. From the rhizobia point of view, several molecules are important in this intimate interaction, highlighting Nod factors, extracellular polysaccharides, and protein secretion systems (López-Baena *et al*., 2016). In case of T3SS, numerous examples have been described to exert neutral, positive or negative effects on symbiosis depending on the plant host (Nelson & Sadowsky, 2015; Jiménez-Guerrero *et al*., 2022). However, studies about the role of rhizobial T6SS on symbiosis are scarce, despite the first report dealing with T6SS indicated that this protein secretion system impairs nodulation between *R. leguminosarum* and pea (Bladergroen *et al*., 2003). Conversely, positive effect of T6SS on symbiosis has been recently reported in the symbiosystems *R. etli-*bean and *Bradyrhizobium-Lupinus* (Salinero-Lanzarote *et al*., 2019, Tighilt *et al*., 2022). In two additional studies, it was demonstrated a role of T6SS on interbacterial competition but not in symbiosis (de Campos *et al*., 2017; Lin *et al*., 2018). In the case of *Sinorhizobium* genera, no studies have been performed so far.

The T6SSs distributed among *Proteobacteria* phylum are classified in five phylogenetic groups (Boyer *et al*., 2009). Reference strains for each phylogenetic group belong to the order *Pseudomonadales*, except for group 5, whose reference strain is *Agrobacterium tumefaciens*. TssB is commonly used to group and classify the different T6SSs among the five phylogenetic groups, since this highly conserved protein conforms the contractile sheath of the T6SS in all microorganisms analyzed so far (Bernal *et al*., 2017; Cascales & Cambillau, 2012). In case of the USDA257, according to its TssB sequence, the T6SS belongs to the group 3, which is one of the most represented phylogenetic groups among the *Rhizobiales* order. An in deep analysis indicates that the genome of USDA257 contains the thirteen genes coding for all structural proteins required for a functional T6SS in one chromosomal cluster. Besides to these T6SS core proteins, *in silico* analysis points out that USDA257 synthetizes two accessory proteins of the membrane complex, TagL and TagM. In addition, most T6SSs described so far are controlled by regulatory proteins (Lin *et al*., 2018; Mougous *et al*., 2007). A well- studied example is the kinase-phosphatase PpkA-PppA couple, which mediates the system formation through a phosphorylation cascade that ends with the structural components assembly. In the case of the USDA257 cluster, we found not only genes coding for PpkA-PppA regulatory proteins but also those encoding the TagF and Fha proteins, which also interact with the system and regulate its activity at post- transcriptional level (Lin *et al*., 2018; Mougous *et al*., 2007). Genes coding effector proteins are often found inside of the T6SS cluster, specifically in the *vgr* gene neighborhood (Durand *et al*., 2014). In USDA257, both *vgrG* and *paar* (*tsre1*) genes encode for evolved proteins. The C-terminal effector domain of VgrG is structurally similar than a polysaccharide lyase-type enzime, which points out that this USDA257 effector could bind to cellulose and polygalacturonic acid, being involved in the attachment to vegetal cell wall. Tsre2 shows structural homology -according to Phyre^2^- with a glycoside hydrolase enzyme with a xyloglucan binding domain able to hydrolyze the plant cell wall (Arnal *et al*., 2019), but also -using Foldseek- with the DNase toxin Tse7 from *P. aeruginosa* PAO1 (Pissaridou *et al*., 2018). This tool also predicts that the Tsre1 protein possesses structural similarity to the antitoxin protein Tse7, further supporting their potential role as a toxin-antitoxin system. Comparisons of this effector across several rhizobial strains indicate that, despite the sequence dissimilarity among the three different versions of the effector, Tsre2 proteins consistently exhibit structural homology with the same two enzymes. This suggests that, regardless of the specific function of this protein, it is likely to be conserved among rhizobia. Moreover, it cannot ruled out that Tsre2 could be a moonlighting effector, a class of multifunctional protein capable of performing more than one distinct biological function, often depending on factors like its location within the cell, interaction with different molecules, or changes in environmental conditions (Jeffery, 1999). It has been recently proposed that moonlighting proteins may be particularly important when the symbiotic interaction becomes more intimate, that is, during the development of the symbiosome (Ma *et al*., 2024).

Overall, the exhaustive *in silico* analysis performed in this work points to a functional T6SS of *S. fredii* USDA257, which might be involved in the interaction with host plant cells and/or with bacteria present in the plant rhizosphere. But is indeed this T6SS fully functional and involved in the direct or indirect optimization of the nodulation process? According to our results, the USDA257 T6SS genes are highly expressed within legume nodules and in poor media at exponential phase, but not in the presence of *nod*-gene inducing flavonoids. Findings described for *R. etli* T6SS are largely similar, since this protein secretion system is fully functional in yeast mannitol medium at high optical densities of growth and in the bean nodules (Salinero-Lanzarote *et al*., 2019). In this bacterium, the T6SS exerts a positive effect in the symbiotic process with its natural legume, whereas information about its role on bacterial competition is missing. The involvement on the symbiotic process of the USDA257 T6SS has been confirmed in nodulation and competitiveness for nodule occupancy assays, where the absence of this protein secretion system reduces the formation of nodules and their relative nodule colonization in soybean. In summary, *in vivo* assays strenghthen, at least in some extend, what emerged from *in silico* analysis, i.e. this protein secretion system is fully functional playing a positive effect on the symbiosis stablished between *S. fredii* USDA257 and *G. max* cv Pekin, its natural host.

So far, the involvement of rhizobial T6SS on interbacterial competition seems to be incompatible for its role on symbiosis and *vicerversa*, since both abilities have never been described at the same time in the same rhizobial strain (Bladergroen *et al*., 2003; de Campos *et al*., 2017; Lin *et al*., 2018; Salinero-Lanzarote *et al*., 2019; Tighilt *et al*., 2022). Competition assays performed against various rhizobacteria under *in vitro* T6SS-inducing conditions indicate that this protein secretion system does not confer killing ability to USDA257. However, this result disagrees with part of the *in silico* analysis of Tsre2 effector, since this protein seems to share structurally homology with a DNase- type toxin (Pissaridou *et al*., 2018). Perhaps the ability to kill other competing rhizobacteria occurs in later stages of symbiosis, during root colonization and infection. This could explain why, under controlled laboratory conditions, we did not detect the toxic effect mediated by Tsre2.

The ecological success of rhizobia is based on their great capacity to adapt, not only to environmental changes as free-living soil bacteria, but also to the challenges they face during root colonization, invasion through the infection tube, and their establishment as mature nitrogen-fixing bacteroids in nodule cells. So far, studies have primarily focused on the characterization of nodulation factors, surface polysaccharides, and the T3SS. However, our results point to a new level of specificity mediated by T6SS effectors, which optimize the symbiotic efficiency by enhancing rhizobial competitiveness during nodulation. Despite possible functions have been suggested for some of these effector proteins, future work is needed to elucidate their functioning and to understand the specific role they play in rhizosphere and symbiosis.

## Supplementary Materials

Figure S1: Southern blot hybridization with probes of *tssA* genes. Figure S2: Multiple alignment of the TssM, TssL, TagM and TagL sequences. Figure S3-5: Multiple alignment of the Tsre1 sequences. Figure S6-8: Multiple alignment of the Tsre2 sequences. Figure S9: Induction of *ppkA* promoter in the presence of *nod* gene inducing-flavonoids. Table S1. Bacterial strains and plasmid used in this study. Table S2. DNA oligonucleotide primers used in this study. Table S3. Nitrogen-fixing and related bacteria harboring a T6SS.

## Acknowledgments and Funding

This work was funded with Projects PID2020- 118279RA-I00 and PID2019-107634RB-I00 funded by MICIU/AEI/10.13039/501100011033.

## Author Contributions

Conceptualization, F.P-M., P.B., and I.J-G.; Methodology, P.R-P., I.J-G., and P.B.; Validation, P.R-P., A.S-R., C.C., J.G., N.M-C., F.P-M, F.O., and I.J-G.; Formal Analysis, P.R-P., F.P-M., P.B. and I.J-G.; Investigation, P.R-P., A.S- R., C.C., N.M-C., F.O., F.P-M, and I.J-G.; Writing – Original Draft Preparation, F.P-M.; Writing – Review & Editing, P.B. and I.J-G.; Visualization, P.R-P., A.S-R., F.P-M., and P.B.; Project Administration, I. J-G, P. B., and F.P-M.; Funding Acquisition, P. B., and F.P-M.; Supervision: I. J-G., P. B., J.G., and F.P-M.

## Conflicts of Interest Statement

The authors declare no conflict of interest.

**Data Availability Statement:** The data are available in the manuscript.

