## Supplementary material for "The functional type VI secretion system of *Sinorhizobium fredii* USDA257 is required for a successful nodulation with *Glycine max* cv Pekin": Suppl Figures

**Supplementary Figures**

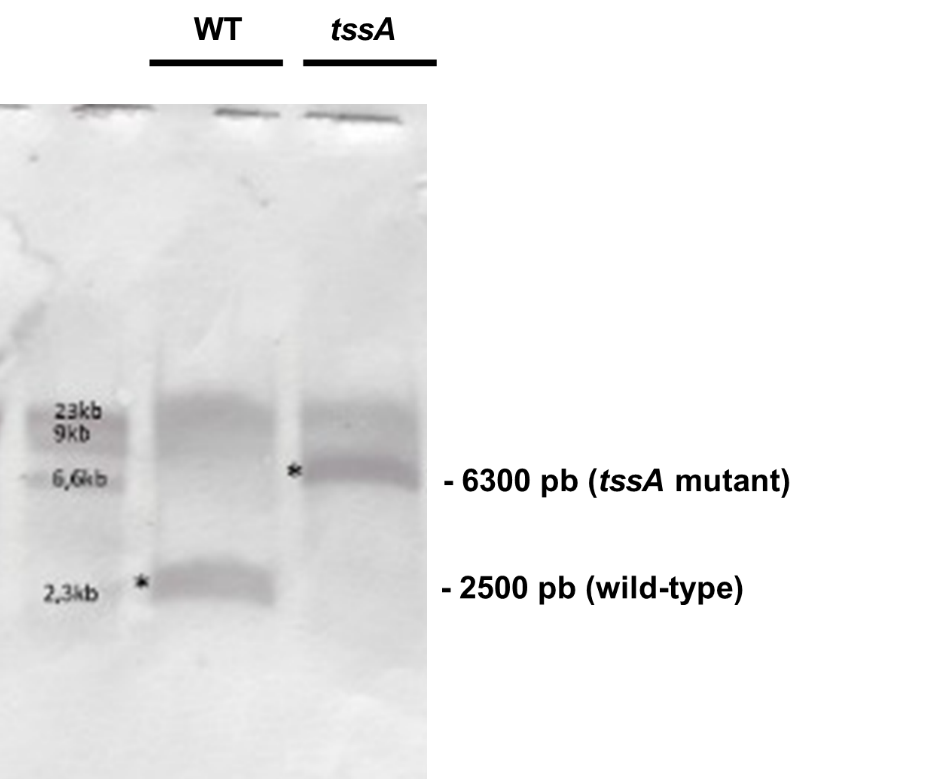

**Figure S1**. Southern blot hybridization using a probe of *tssA* gene (made with primers listed in Table S2). The gDNA of each USDA257 strain was digested with *Kpn*I restriction enzyme.

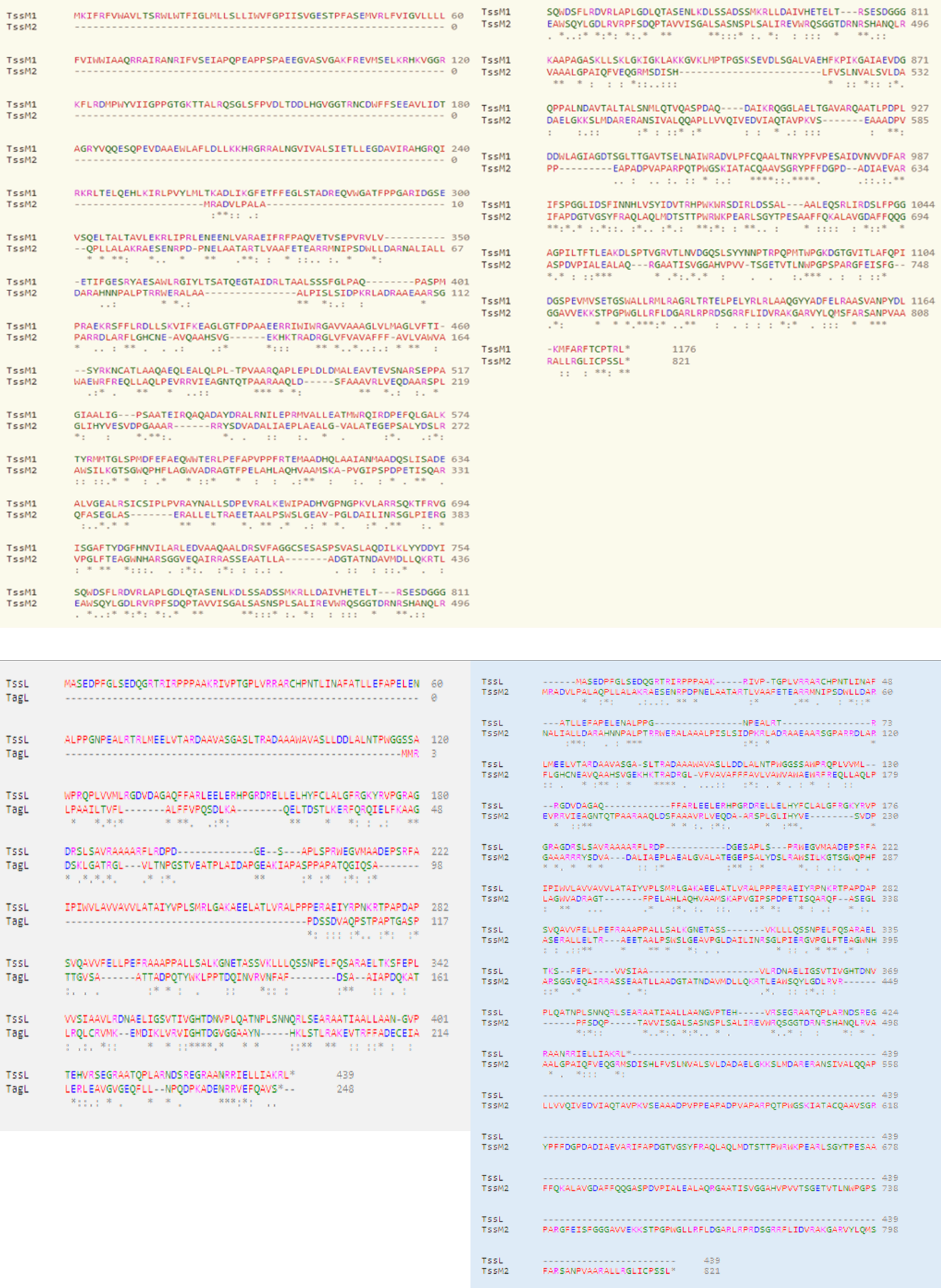

**Figure S2**. Multiple alignment of the TssM, TssL, TagM and TagL sequences. Representation of alignment of the protein sequences of TssM and TagM (TssM2) (green), of the sequences of TssL and TagL (grey) and of the sequences of TssL and TagM (TssM2) (blue).

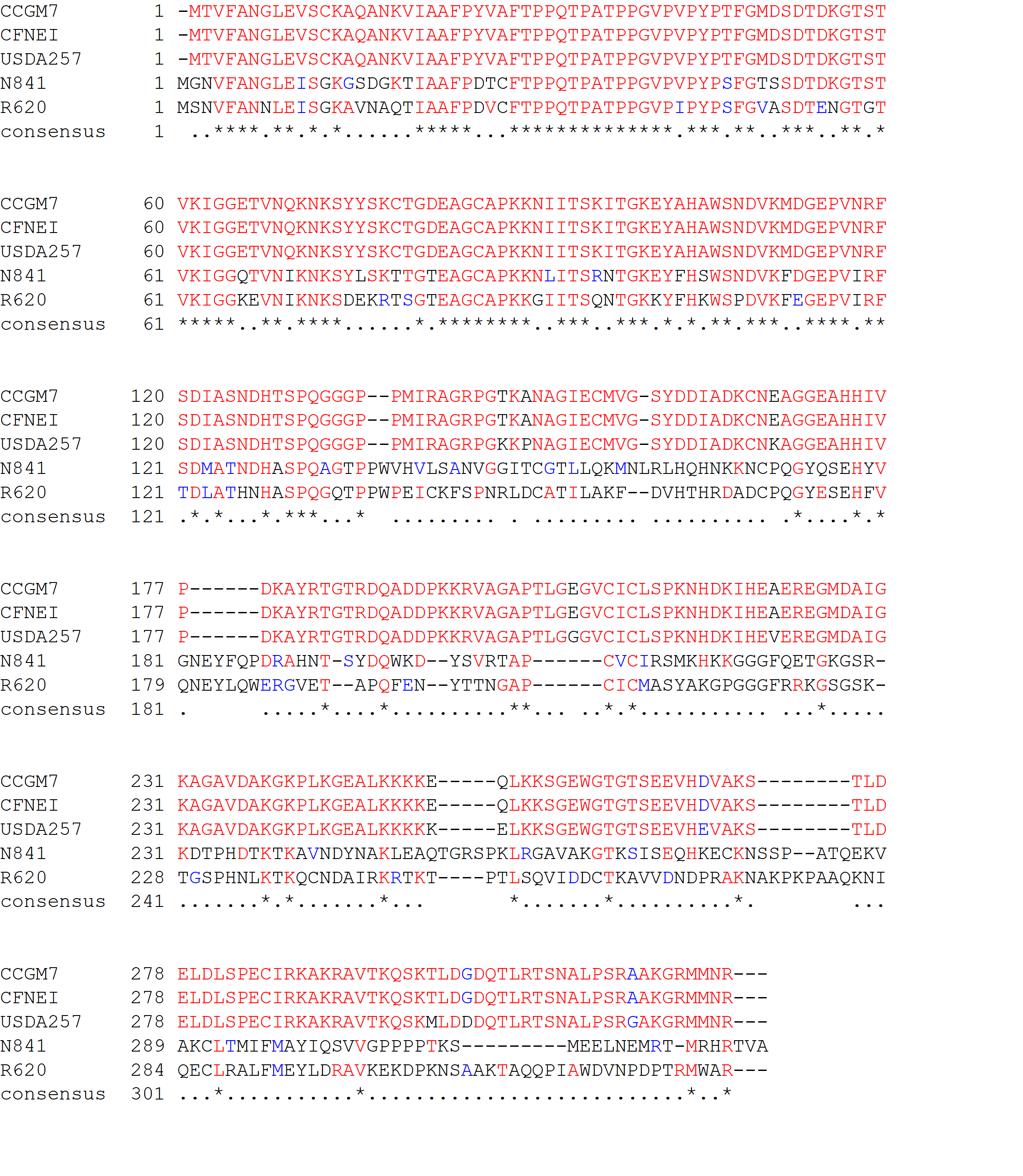

**Figure S3**. Multiple alignment of the Tsre1 sequences. Representation of the Tsre1 protein alignment from *S. fredii* USDA257, *Sinorhizobium americanum* strains CCGM7 and CFNEI 73. *Rhizobium phaseoli* strains N841 and N620 are included for comparative purposes.

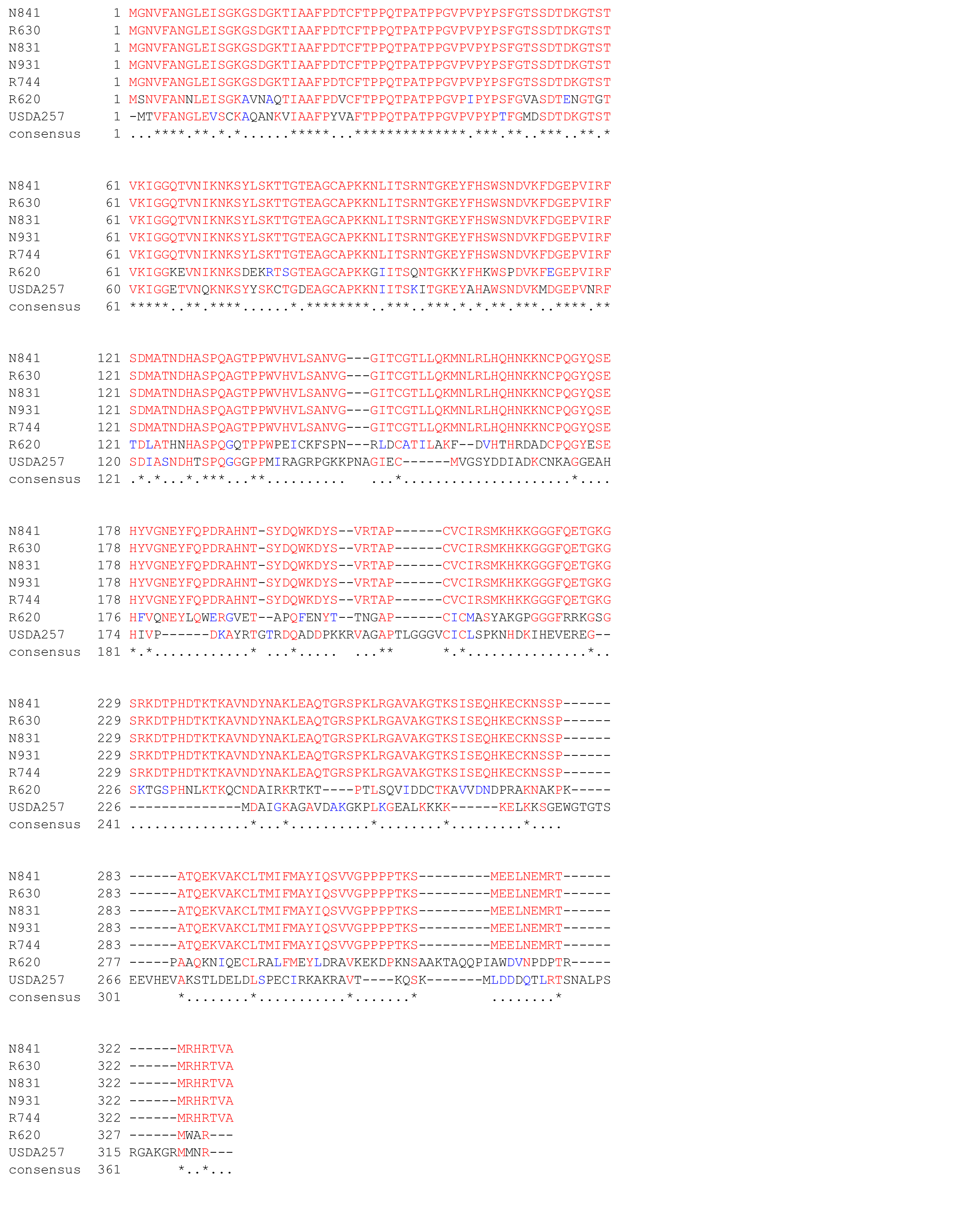

**Figure S4**. Multiple alignment of the Tsre1 sequences. Representation of the Tsre1 protein alignment from *Rhizobium phaseoli* strains N841, R630, N831, N931 and R744. *S***.***fredii* USDA257 and *Rhizobium phaseoli* N620 are included for comparative purposes.

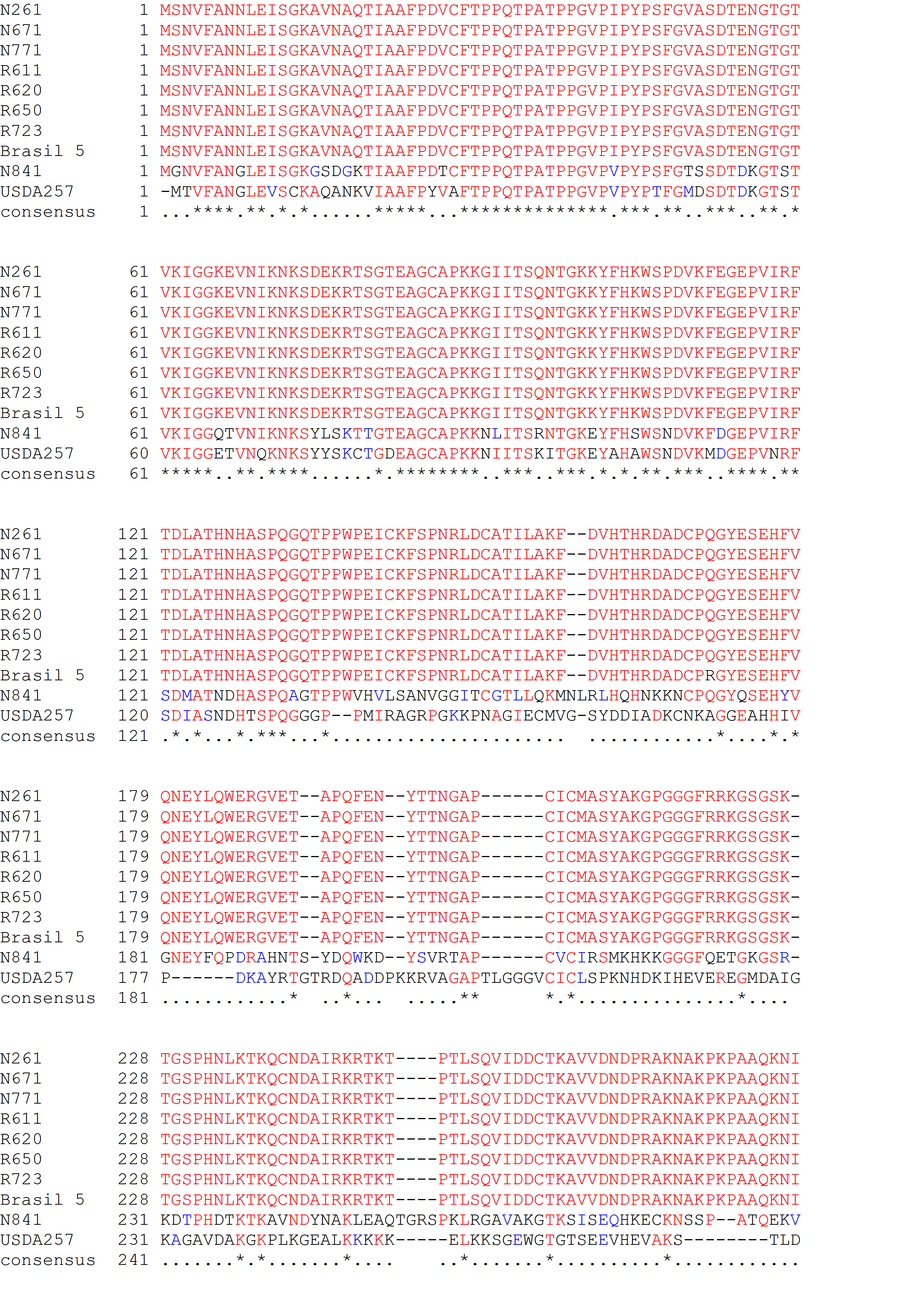

**Figure S5**. Multiple alignment of the Tsre1 sequences. Representation of the Tsre1 protein alignment from *Rhizobium phaseoli* strains R620, R650, R611, N771, N671, N261, R723 and Brasil 5. *S. fredii* USDA257 and *Rhizobium phaseoli* N841 are included for comparative purposes.

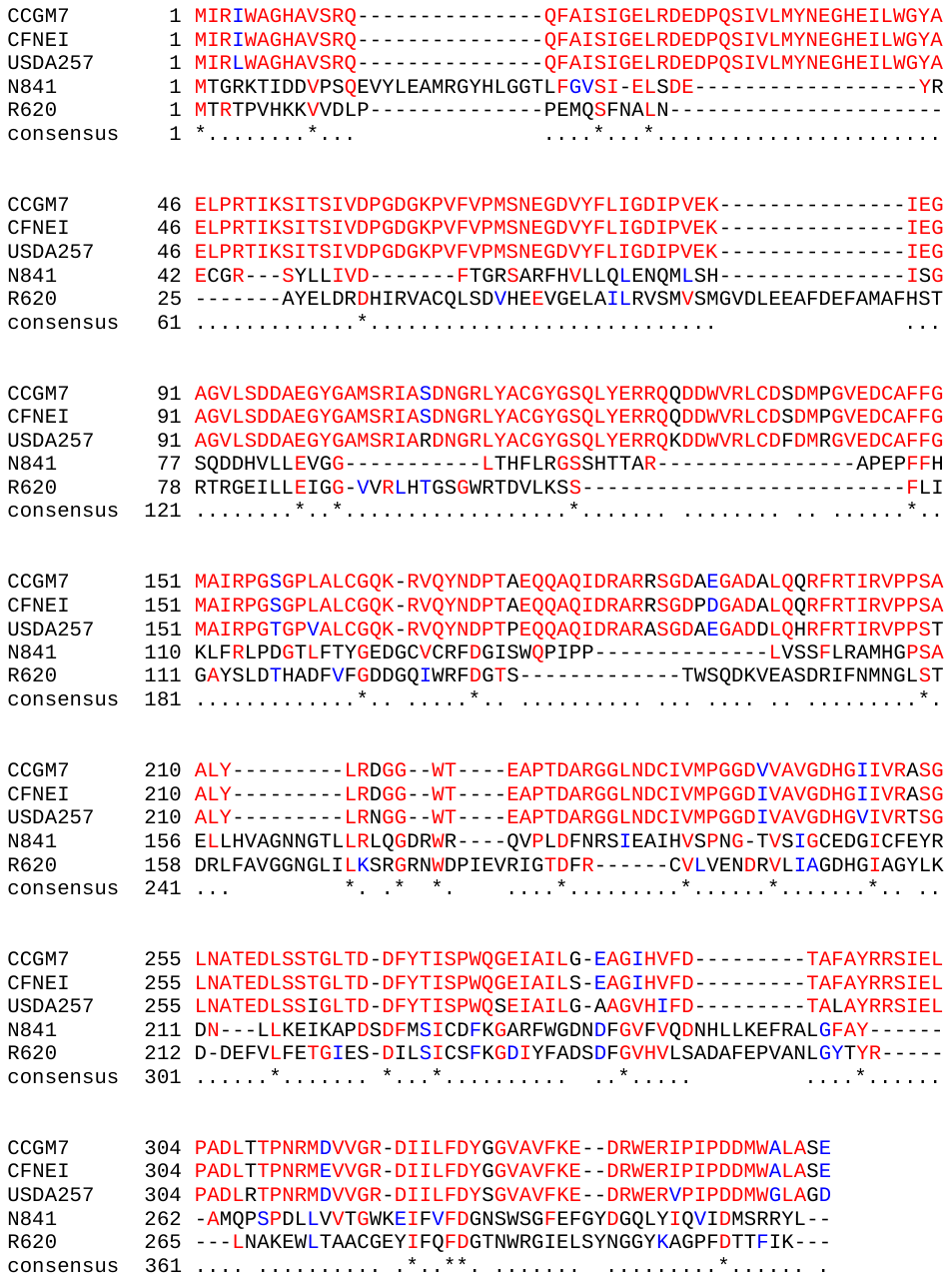
**Figure S6**. Multiple alignment of the Tsre2 sequences. Representation of the Tsre2 protein alignment from *S*. *fredii* USDA257, *Sinorhizobium americanum* strains CCGM7 and CFNEI 73. *Rhizobium phaseoli* strains N841 and N620 are included for comparative purposes.

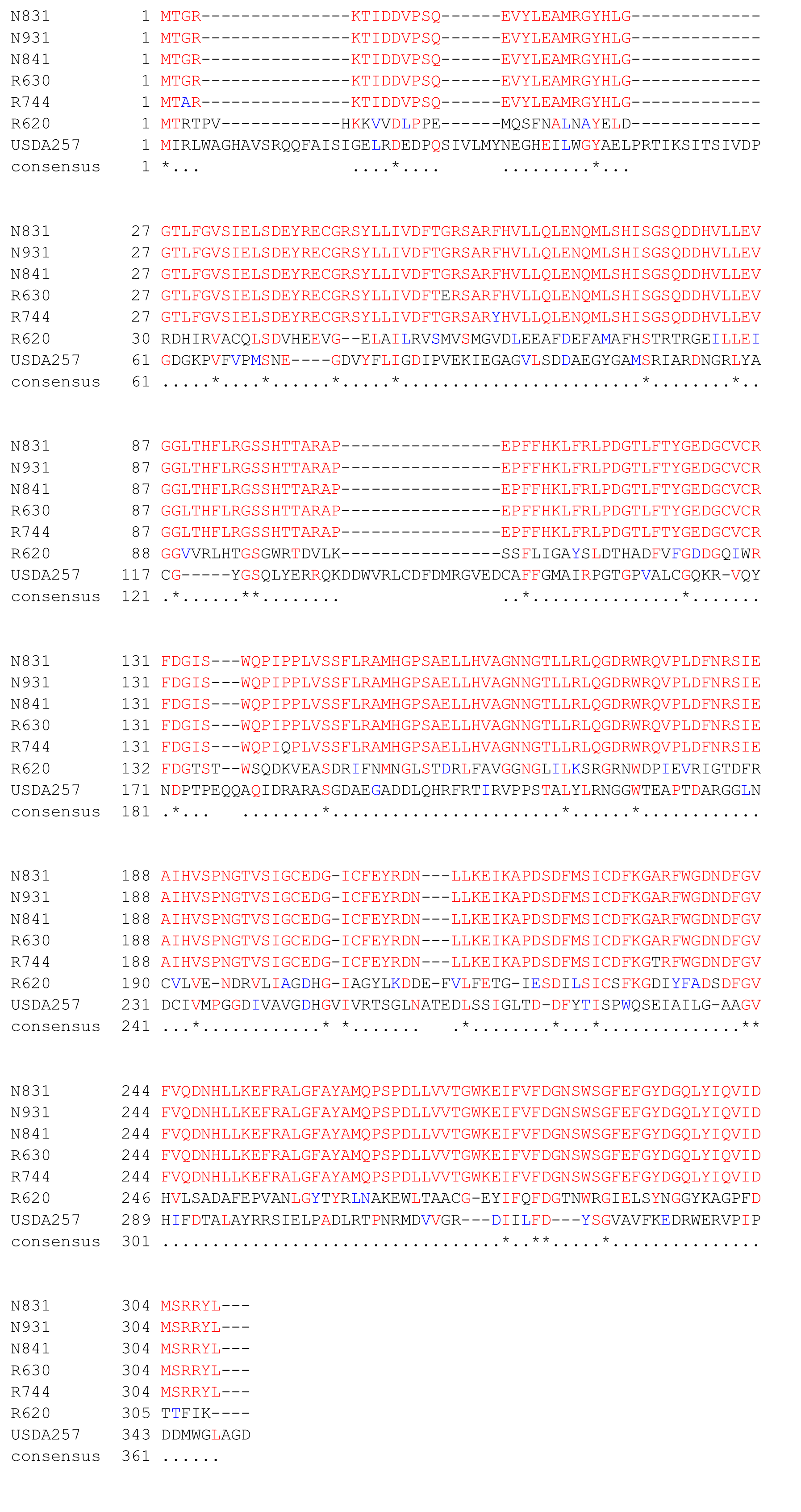

**Figure S7**. Multiple alignment of the Tsre2 sequences. Representation of the Tsre2 protein alignment from *Rhizobium phaseoli* strains N841, R630, N831, N931 and R744. *S. fredii* USDA257 and *Rhizobium phaseoli* N620 are included for comparative purposes.

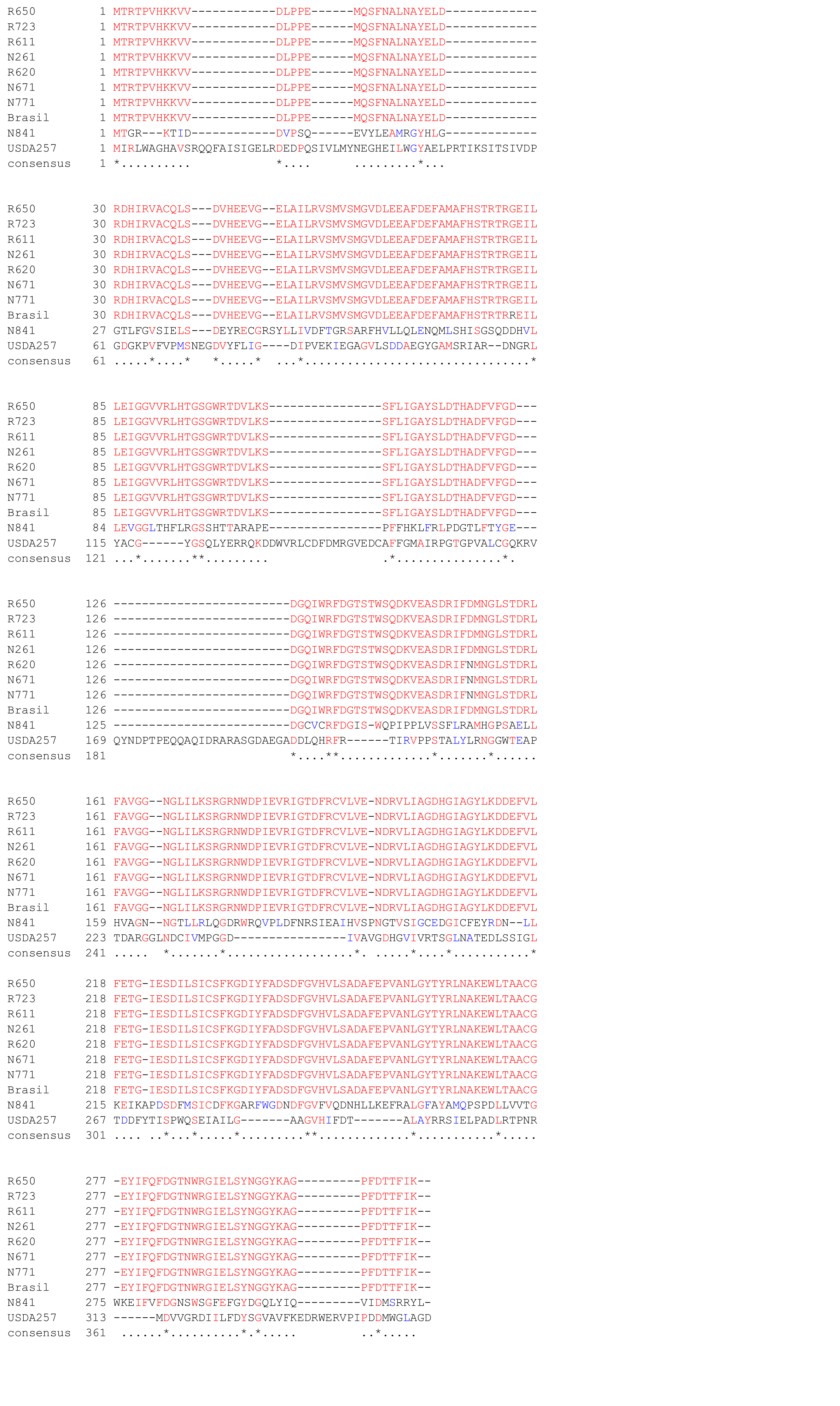

**Figure S8**. Multiple alignment of the Tsre2 protein sequences. Representation of the Tsre2 protein alignment from *Rhizobium phaseoli* strains R620, R650, R611, N771, N671, N261, R723 and Brasil 5. *S. fredii* USDA257 and *Rhizobium phaseoli* N841 are included for comparative purposes.

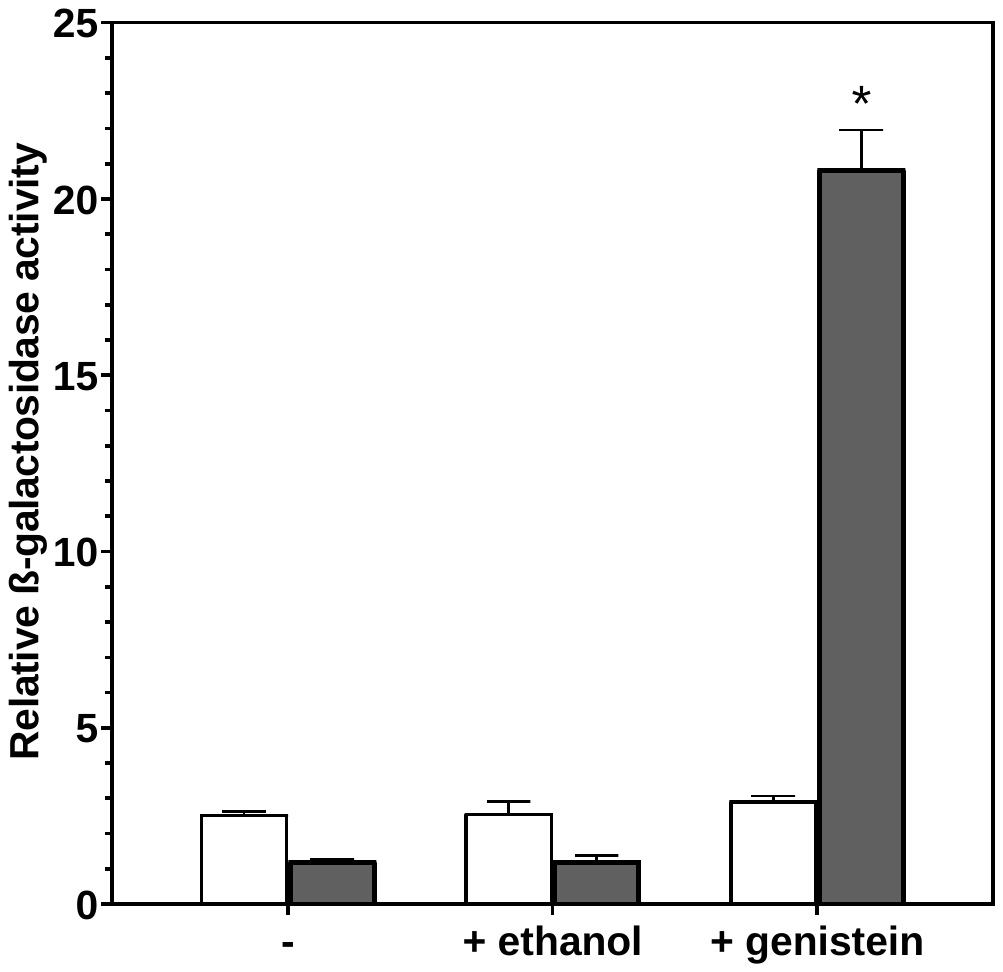

**Figure S9**. Fold-change values of the β-galactosidase activities of *S. fredii* USDA257 carrying plasmid pMP220 containing the *ppkA* (white bars) promoter fused to the *lacZ* gene, or plasmid pMP240, which contains the conserved *nodA* promoter from *R. leguminosarum* fused to *lacZ* (dark grey bars). Assayed conditions were MM3 medium at stationary phase in the absence and in the presence of ethanol or the USDA257-inducing flavonoid genistein. Parameters were individually compared with those obtained by in the wild-type strain carrying the empty vector in MM3 medium by using the t test with multiple comparisons. Values tagged by asterisks (*) are significantly different at the level of α = 5%.
