## Supplementary material for "The functional type VI secretion system of *Sinorhizobium fredii* USDA257 is required for a successful nodulation with *Glycine max* cv Pekin": Suppl Tables

**Supplementary Tables**

**Table S1.** Bacterial strains and plasmid used in this study.

| **Strain or plasmid** | **Relevant properties** | **Source or reference** |
| --- | --- | --- |
| ***Sinorhizobium fredii*** | | |
| USDA257 | Wild-type strain, Rif^R^ | (Pueppke & Broughton, 1999) |
| USDA257 *tssA* | USDA257 *tssA*::pK18*mob*, Rif^R^ Km^R^ | This work |
| USDA257 (pMP220::P_tssA_) | Wild-type strain carrying plasmid pMP220::P_tssA_; Rif^R^, Tc^R^ | This work |
| USDA257 (pMP220::P_ppkA_) | Wild-type strain carrying plasmid pMP220::P_ppkA_; Rif^R^, Tc^R^ | This work |
| USDA257 (pMP240) | Wild-type strain carrying plasmid pMP240; Rif^R^, Tc^R^ | This work |
| USDA257 (pMP220) | Wild-type strain carrying plasmid pMP220; Rif^R^, Tc^R^ | This work |
| USDA257  (pBBR4::P_kan_::*gfp*-P_ppkA_::*rfp*) | Wild-type strain carrying plasmid pBBR4::P_kan_::*gfp*-P_ppkA_::*rfp*; Rif^R^, Ap^R^ | This work |
| USDA257 (pBBR4::P_kan_::*gfp*-P_w/o_::*rfp*) | Wild-type strain carrying plasmid pBBR4::P_kan_::*gfp*-P_w/o_::*rfp*; Rif^R^, Ap^R^ | This work |
| HH103 | Wild-type strain, Str^R^ | (Buendía-Clavería et al., 1989) |
| ***Agrobacterium tumefaciens*** | | |
| C58 (pRL662::*gfp2*) | Wild-type strain carrying the GFP fluorescent marker, Gm^R^ | Bernal et al., 2017 |
| ***Pectobacterium carotovorum*** | | |
| SCRI 194 (pRL662::*gfp2*) | Wild-type strain carrying the GFP fluorescent marker, Gm^R^ | Bernal et al., 2017 |
| ***Escherichia coli*** | | |
| DH5α | *supE44 ΔlacU169 hsdR17 racA1 endA1 gyr96 thi-1 relA1* | Stratagene (USA) |
| DB3.1 | *gyrA462*, *endA1*, ∆(*sr1-recA*), *mcrB*, *mrr*, *hsdS20*, *glnV44* (=*supE44*), *ara14*, *galK2*, *lacY1*, *proA2*, *rpsL20*, *xyl5*, *leuB6*, *mtl1* | Invitrogen (USA) |
| BL21(DE3) | *E. coli* B F^-^*dcmompThsdS* (r_B_^-^m_B_^-^) *gallon*λ (DE3 [*lacIlac*UV5-T7 gene *1ind1sam7nin5*]) | Studier and Moffatt, 1986 |
| **Plasmids** | | |
| pRK2013 | Helper plasmid, Km^R^ | Figurski & Helinski, 1979 |
| pRL662::*gfp2* | Plasmid expressing the GFP2, Gm^R^ | Bernal et al., 2017 |
| pK18*mob* | Cloning vector, suicide in rhizobia, Km^R^ | Schäfer et al., 1994 |
| pMP220 | Cloning vector, empty plasmid, Tc^R^ | Spaink et al., 1987 |
| pMP240 | pMP220 plasmid carrying the transcriptional fusion between the *R. leguminosarum* bv. *viciae nodA* promoter and the *lacZ* gene, Tc^R^ | de Maagd et al., 1988 |
| pMUS1480 | pK18*mob* plasmid carrying an internal fragment of the USDA257 *tssA* gene, Km^R^ | This work |
| pMP220::P_tssA_ | pMP220 plasmid carrying the transcriptional fusion between the USDA257 *hcp* promoter region and the *lacZ* gene, Tc^R^ | This work |
| pMP220::P_ppkA_ | pMP220 plasmid carrying the transcriptional fusion between the USDA257 *tssA* promoter region and the *lacZ* gene, Tc^R^ | This work |
| pBBR4::P_kan_::*gfp*-P_w/o_::*rfp* | pBBR4 plasmid carrying the transcriptional fusion between the kanamycin gene promoter (constitutive) and the *gfp* gene, Ap^R^ | Samal and Chatterjee, 2021 |
| pBBR4::P_kan_::*gfp*-P_ppkA_::*rfp* | pBBR4 plasmid carrying the transcriptional fusions between the kanamycin gene promoter (constitutive) with the *gfp* gene and the *ppkA* promoter (T6SS-controlled) with the *rfp* gene, Ap^R^ | This work |
| pDONR207 | Gateway donor vector; Gm^R^ | Invitrogen (USA) |
| pET42 | Plasmid with His N and C terminal tag for bacterial expression; Ap^R^ | Novagen (Germany) |
| pDONR207::*hcp* | pDONR207 carrying the USDA257 *hcp* ORF tagged with His in both terminal ends; Gm^R^ | This work |
| pET42::*hcp* | pET28a carrying the USDA257 *hcp* ORF tagged with His in both terminal ends; used for antibody obtention; Ap^R^ | This work |

* Ap^R^, Gm^R^, Km^R^. Tc^R^ and Rif^R^ indicate resistance to ampicillin, gentamicin, kanamycin, tetracycline and rifampicin, respectively.

### Table S2. DNA oligonucleotide primers used in this study.

| Name | Sequence | Usage |
| --- | --- | --- |
| tssA_ecoRI-F | 5'- ATAGAATTCATGGGCGATTCCACGAAC | Amplification of an internal fragment of *tssA* gene |
| tssA_bamHI-R | 5'- ATAGGATCCGTCGCCCAATAACCCATC |  |
| PppkAecoRI-F | 5'- ATTGAATTCTGGATGGCGTGAAAGATGTG | Amplification of the promoter region of *ppkA* gene |
| PppkAecoRI-R | 5'- ATTGAATTCGTAGGTGTTGTTGAGCACCT |  |
| rfppromExt-F | 5´-CTTGGAGCCGTACTGGAAC | Sequence verification of inserts cloned into pBBR4::P_kan_::*gfp*-P_w/o_::*rfp* |
| rfppromExt-R | 5´-CAGCACGTGTCTTGTAGTTCC |  |
| PtssA_ecoRI-F | 5'- ATATGAATTCACGGCGACAAGTCCGAAC | Amplification of the promoter region of *ttsA* gene |
| PtssA_xbaI-R | 5'- ATAATCTAGATCGAACAACAGATCCCCCC |  |
| PppkA-ecoRI-F | 5'- ATATGAATTCACCTCTTCGGATCCGAACTG | Amplification of the promoter region of *ppkA* gene |
| PppkA-xbaI-R | 5'- ATAATCTAGAGTAGGTGTTGTTGAGCACCT |  |
| hcp_attB1 | 5'- GGGGACAAGTTTGTACAAAAAAGCAGGCTTAATGAAAATTGATGGATTTCTAAA -3 | Used for cloning of the USDA257 *hcp* ORF without the stop codon in plasmid pDONR207 by the Gateway system |
| hcp_attB2_ns | 5'-GGGGACCACTTTGTACAAGAAAGCTGGGT-3 |  |
| pDONR_F | 5'-CGTTAACGCTAGCATGGATCTC-3' | Sequence verification of inserts cloned into pDONR207 |
| pDONR_R | 5'-GTAACATCAGAGATTTTGAGAC-3' |  |
| qtssA-F | 5'- TGCTGAATTCCTCGGAAG | *q*PCR assays |
| qtssA-R | 5'- CAGCATCGACTTGACGAA |  |
| q16S-F | 5'- TAAACCACATGCTCCACC | *q*PCR assays |
| q16S-R | 5'- GATACCCTGGTAGTCCAC |  |

**Table S3.** Nitrogen-fixing and related bacteria harboring a T6SS.

| **Order Rhizobiales**  **(α-Proteobacteria)** | **Family** | **Genus** | **Species** | **Strains number** | **Function** | **References** |
| --- | --- | --- | --- | --- | --- | --- |
|  | *Brucellaceae* |  |  |  |  |  |
|  |  | *Brucella* (*Ochrobactrum)* |  |  |  |  |
|  | *Bradyrhizobiaceae (Nitrobacteraceae)* |  |  |  |  |  |
|  |  | *Bradyrhizobium* |  |  |  |  |
|  |  |  | *japonicum* | 1 |  |  |
|  |  |  | *diazoefficiens* | 2 |  |  |
|  |  |  | *elkanii* | 1 |  |  |
|  |  |  | *canariense* | 1 |  |  |
|  |  |  | *pachyrhizi* | 1 |  |  |
|  |  |  | *vignae* | 0 |  |  |
|  |  | *Blastobacter* |  | 0 |  |  |
|  |  |  | *denitrificans* | 0 |  |  |
|  | *Hyphomicrobiaceae* |  |  |  |  |  |
|  | [*Devosiaceae*](https://en.wikipedia.org/wiki/Devosiaceae) |  |  |  |  |  |
|  |  | *Devosia* |  |  |  |  |
|  |  |  | *neptuniae* | 0 |  |  |
|  |  |  | *sp Root635* | 1 |  |  |
|  | *Methylobacteriaceae* |  |  |  |  |  |
|  |  | *Microvirga* |  | 2 |  |  |
|  |  | *Methylobacterium* |  | 21 |  |  |
|  |  |  | *sp* | 12 |  |  |
|  |  |  | *segetis* | 1 |  |  |
|  |  |  | *organophilum* | 1 |  |  |
|  |  |  | *platani* | 1 |  |  |
|  |  |  | *crusticola* | 1 |  |  |
|  |  |  | *indicum* | 1 |  |  |
|  |  |  | *nodulans* | 1 |  |  |
|  |  |  | *variabile* | 1 |  |  |
|  |  |  | *tarhaniae* | 1 |  |  |
|  |  |  | *dankookense* | 1 |  |  |
|  | *Methylocystaceae* |  |  |  |  |  |
|  |  | *Methylocystis* |  |  |  |  |
|  |  |  | *heyeri* | 1 |  |  |
|  |  | *Methylosinus* | *sp.* | 4 |  |  |
|  | *Phyllobacteriaceae* |  |  |  |  |  |
|  |  | *Mesorhizobium* |  | 17 |  |  |
|  |  |  | *loti* | 1 |  |  |
|  |  |  | *mediterraneum* |  |  |  |
|  |  |  | *tianshanense* |  |  |  |
|  |  |  | *ciceri* | 1 |  |  |
|  |  |  | *qingshengii* | 1 |  |  |
|  |  |  | *zhangyense* | 1 |  |  |
|  |  |  | *amorphae* | 1 |  |  |
|  |  |  | *ephedrae* | 1 |  |  |
|  |  |  | *alhagi* | 1 |  |  |
|  |  |  | *waimense* | 1 |  |  |
|  |  |  | *huakuii* | 1 |  |  |
|  |  |  | *temperatum* | 1 |  |  |
|  |  |  | *sp.* | 7 |  |  |
|  |  | *Aminobacter* |  | 4 |  |  |
|  |  |  | *aminovorans* | 1 |  |  |
|  |  |  | *sp.* | 3 |  |  |
|  |  | *Phyllobacterium* |  | 5 |  |  |
|  |  |  | *myrsinacearum* | 1 |  |  |
|  |  |  | *phragmitis* | 1 |  |  |
|  |  |  | *leguminum* | 1 |  |  |
|  |  |  | *sp.* | 2 |  |  |
|  | *Rhizobiaceae* |  |  |  |  |  |
|  |  | *Agrobacterium* |  | 19 |  |  |
|  |  |  | *tumefaciens* | 5 |  |  |
|  |  |  | *larrymoorei* | 1 |  |  |
|  |  |  | *bohemicum* | 1 |  |  |
|  |  |  | *rosae* | 1 |  |  |
|  |  |  | *vitis* | 1 |  |  |
|  |  |  | *rhizogenes* | 1 |  |  |
|  |  |  | *rubi* | 1 |  |  |
|  |  |  | *deltaense* | 1 |  |  |
|  |  |  | *radiobacter* | 1 |  |  |
|  |  |  | *genomosp* | 1 |  |  |
|  |  |  | *salinitolerans* | 1 |  |  |
|  |  |  | *sp* | 4 |  |  |
|  |  | *Ensifer* |  | 3 |  |  |
|  |  |  | *glycinis* | 1 |  |  |
|  |  |  | *sp* | 2 |  |  |
|  |  | *Sinorhizobium* (*=Ensifer*) |  | 6 |  |  |
|  |  |  | *fredii* | 1 |  |  |
|  |  |  | *americanum* | 1 |  |  |
|  |  |  | *saheli* | 2 |  |  |
|  |  |  | *medicae* | 1 |  |  |
|  |  |  | *sp* | 1 |  |  |
|  |  |  | *meliloti* |  |  |  |
|  |  |  | *terangae* |  |  |  |
|  |  |  | *xinjiangense* |  |  |  |
|  |  | *Allorhizobium* |  |  |  |  |
|  |  |  | *oryzae* | 0 |  |  |
|  |  |  | *terrae* | 1 |  |  |
|  |  | *Shinella* |  |  |  |  |
|  |  |  | *kummerowiae* | 0 |  |  |
|  |  |  | *fusca* | 1 |  |  |
|  |  | *Neorhizobium* |  |  |  |  |
|  |  |  | *galegae* | 1 |  |  |
|  |  | *Pararhizobium* |  | 0 |  |  |
|  |  |  | *giardinii* | 0 |  |  |
|  |  | *Hoeflea* | *sp* | 1 |  |  |
|  |  | *Rhizobium* |  | 28 |  |  |
|  |  |  | *etli* | 1 |  |  |
|  |  |  | *etli* bv*. mimosae* Mim1 | 1 | Nodulation | Salinero-Lanzarote et al., 2019 |
|  |  |  | *acidisoli* | 1 |  |  |
|  |  |  | *aegyptiacum* | 1 |  |  |
|  |  |  | *phaseoli* | 1 |  |  |
|  |  |  | *tropici* | 1 |  |  |
|  |  |  | *leguminosarum* bv*. viciae* |  |  |  |
|  |  |  | *leguminosarum* bv*. trifolii* | 1 | Nodulation | Bladergroen et al., 2003 |
|  |  |  | *leguminosarum* bv*. phaseoli* CCGM1 | 1 |  |  |
|  |  |  | *sullae* |  |  |  |
|  |  |  | *leucaneae* |  |  |  |
|  |  |  | *album* | 1 |  |  |
|  |  |  | *deserti* | 1 |  |  |
|  |  |  | *oryziradicis* | 1 |  |  |
|  |  |  | *sophoriradicis* | 1 |  |  |
|  |  |  | *hidalgonense* | 1 |  |  |
|  |  |  | *esperanzae* | 1 |  |  |
|  |  |  | *taibaishanense* | 1 |  |  |
|  |  |  | *gallicum* | 1 |  |  |
|  |  |  | *giardinii* |  |  |  |
|  |  |  | *galegae* |  |  |  |
|  |  |  | *cellulosilyticum* |  |  |  |
|  |  |  | *lupini* |  |  |  |
|  |  |  | *lusitanum* |  |  |  |
|  |  |  | *mongolense* |  |  |  |
|  |  |  | *sp* | 13 |  |  |
|  | *Xanthobacterecaceae* |  |  |  |  |  |
|  |  | *Azorhizobium* |  |  |  |  |
|  |  |  | *caulinodans ORS571* | 1 | Interbacterial competition | Lin et al., 2018 |
|  |  |  | *oxalatiphilum* | 1 |  |  |
| **Order Burkholderiales Order**  **(β-Proteobacteria)** | **Family** | **Genus** | **Species** | **Strains number** | **Function** | **References** |
|  | *Burkholderiaceae* |  |  |  |  |  |
|  |  | *Burkholderia* |  |  |  |  |
|  |  |  | *pyrrocinia* | 1 |  |  |
|  |  | *Paraburkholderia* |  |  |  |  |
|  |  |  | *tuberum* | 1 |  |  |
|  |  |  | *phymatum* |  |  |  |
|  |  |  | STM815 | 1 (2 clusters) | Nodulation | Hug et al., 2021 |
|  |  |  | LMG 21445T | 0 | Interbacterial competition | de Campos et al., 2017 |
|  |  |  | *mimosarum* | 1 |  |  |
|  |  |  | *tropica* | 1 |  |  |
|  |  |  | *graminis* C4D1M | 1 (2 clusters) |  |  |
|  |  | *Cupriavidus* |  |  |  |  |
|  |  |  | *taiwanensis* LMG19424 | 1 |  |  |
|  |  |  | *necator* | 1 |  |  |
| **Others** | **Family** | **Genus** | **Species** | **Strains number** | **Function** | **References** |
|  |  | *Bacteroides* |  | 1 |  |  |
|  |  |  | *fragilis* |  |  |  |
|  |  | *Pseudomonas* |  | 2 |  |  |
|  |  |  | *putida* | 2 |  |  |
|  |  |  | *aeruginosa* | 1 |  |  |
